# A repertoire of foraging decision variables in the mouse brain

**DOI:** 10.1101/2021.04.01.438090

**Authors:** Fanny Cazettes, Luca Mazzucato, Masayoshi Murakami, Joao P. Morais, Alfonso Renart, Zachary F. Mainen

## Abstract

In any given situation, the environment can be parsed in different ways to define useful decision variables (DVs) for any task, but the way in which this manifold of potential decision strategies is processed to shape behavioral policies is not known. We recorded neural ensembles in the frontal cortex of mice performing a foraging task admitting multiple DVs. Methods developed to uncover the currently employed DV revealed the use of multiple strategies and latent changes in strategy within sessions. Optogenetic manipulations showed that the secondary motor cortex (M2) is needed for mice to use the different DVs in the task. Surprisingly, we found that, regardless of the DV best explaining the behavior of each mouse, M2 activity reflected a full basis set of computations spanning a repertoire of DVs extending beyond those useful for the present task. This form of multiplexing may confer considerable advantages for learning and adaptive behavior.

## INTRODUCTION

An adaptive strategy to control behavior is to take actions that lead to good outcomes given that the environment is in a particular state. Yet, environmental states are often complex, with manifold sources of potentially relevant information, some that are directly observable and others that can only be revealed through a process of inference. Therefore, an agent typically also faces the problem of selecting which are the environmental variables on which to base a decision and how must these variables be processed algorithmically to reveal the appropriate ‘decision variable’ (DV). The problem of selecting a DV is likely a more difficult computational problem faced by a decision maker than the decision itself, but how it is accomplished has received scant investigation^1^.

An important possibility is that an agent need not commit to a particular DV but may entertain several in parallel. The ability to parallelize operations of decision processing, such as temporal integration, would permit adaptation to changes in task contingencies without implementation of new computations, and could therefore potentially speed learning and provide flexibility in combining and switching of strategies. However, little is known about the limitations and possibilities for multiplexing the algorithms used to derive DVs from sensory evidence. On the one hand, behavioral studies in humans suggested that two streams of sensory evidence can only be incorporated into a DV one at a time, necessitating serial processing^2–4^. On the other hand, it has been shown that there exist neurons integrating evidence about a single sensory event with diverse timescales^5^, and that diverse time-scales are present in neurons within local circuits^6^, which could reflect a simple form of algorithmic multiplexing. It thus remains unclear whether various computations can be carried out in parallel on different streams of evidence to form a broad range of simultaneously available DVs.

To directly study the possibility of multiplexing computations on sequential inputs in the brain, we leveraged a foraging task based on processing a stream of binary outcomes (successful and unsuccessful foraging attempts) to inform a decision of whether to leave or stay^7,8^. This task admits multiple strategies for processing the series of outcomes which are associated with different precisely quantifiable DVs. Evaluation of these DVs allows the experimenter to infer the implementation of ‘counterfactual’ strategies, i.e., strategies which are potentially applicable, but unused. This can be done because the task involves precise sequences of stimuli that can be processed in multiple ways. If such counterfactual strategies could be decoded from the brain, it would be evidence for parallel processing of serial information.

Here, using population recordings and optogenetic silencing in the frontal cortex of mice performing the foraging task, we identified a brain region (the secondary motor cortex) where the multiple DVs used by the mice could be decoded simultaneously. Critically, we found that the extent to which each DV was represented in the cortex did not depend on the particular strategy used by each mouse. These observations suggest that mice use an algorithm for decision-making that relies on the parallel computation of multiple DVs in the frontal cortex.

## RESULTS

### Multiple DVs predict switching decision

In our task, a head-fixed mouse collected rewards at a virtual foraging site by licking from a spout (Fig. 1a; Extended Data Fig. 1). During a foraging bout, either a fixed amount of reward (1μL consumed in a single lick) or nothing was delivered for each detected lick. At any time, the mouse could choose to continue licking or give up and explore a new site by starting to run. In particular, there were two virtual foraging sites only one of which was active at a given time and would deliver reward with a probability of 0.9 after each lick. The active site had also a probability of 0.3 of switching after each lick^8^. This switch from active to inactive only happened once while the mouse was at the site, so if it left the site before the switch, no rewards were delivered at the other site (and it had to return to the original site and restart licking). Therefore, the best strategy to time the switching decision was to infer the latent state corresponding to which site was currently active^8^. This inference-based strategy was supported by a particular DV that consisted of temporally accumulating consecutive failures with a complete reset upon receiving a reward (Fig. 1b). This is because a failure to receive reward provides evidence that the active state had switched, whereas a reward always signaled the active state with certainty. Using this strategy, mice would leave the current site when the *‘consecutive failures’* DV reaches a given threshold^8^. Yet, in principle, mice could time their decision to leave by using any number of alternative strategies based on the sequence of rewarded and unrewarded licks regardless of the true causal structure of the task. In fact, early on during training when learning the task, mice do not appear to calculate the inference-based DV^8^. Their behavior is better described by a strategy which does not contemplate discrete transitions to a fully depleted state, and instead relies on a running estimate of the *‘value’* of the current site based on the difference between recently observed rewards and failures (Fig. 1c). Using this strategy, mice decide to abandon a foraging site when its value is sufficiently low (or its negative sufficiently high). We refer to this as a *stimulus-bound strategy* because it treats observable outcomes (the stimuli) as direct – although probabilistic – reporters of the valence of current environmental states, without further assumptions or models about environmental dynamics. For our present purposes, the essential aspect of these two strategies is that they use the same observable outcomes (series of rewarded and unrewarded licks) in qualitatively different ways to update their corresponding DV – a full reset versus a quantitative incremental increase in current value. This allows us to unambiguously identify the two DVs, their behavioral consequences, and their neural representations.

**Figure 1:**
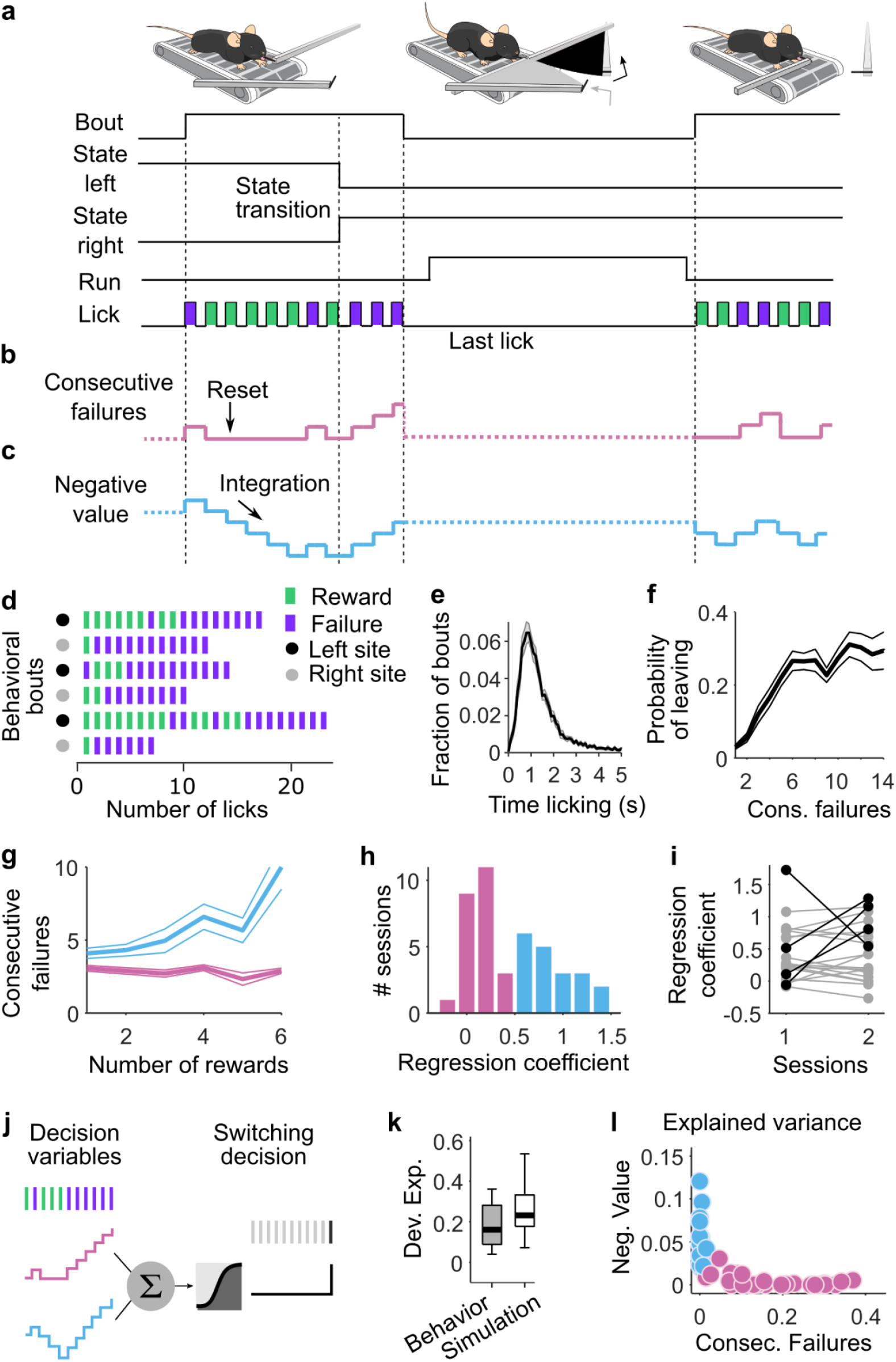
Multiple DVs predict foraging behavior. (a) A head-fixed mouse placed on a treadmill chooses to exploit one of the two foraging sites (two movable arms on each side of the treadmill). A bout of behavior consists of a series of rewarded and unrewarded licks at one of the sites. When a site is in an active state, the probability of each lick being rewarded is 90% and each lick is associated with a 30% probability of state transition. Independently from state transition, animals can choose to switch between sites at any time by running a set distance on the treadmill. During site-switching, the spout in front moves away and the distal one moves into place. (b) The DV that the mouse needs to compute to infer the hidden state of the resource site. (c) Alternative DV supporting a stimulus-bound strategy: the ‘negative value’. (d) Example sequences of observable events during different behavior bouts. (e) Histogram of bout duration (mean ± s.e.m. across sessions; n = 42). (f) Probability of leaving the foraging site as a function of the number of consecutive failures after the last reward (mean ± s.d. across mice). (g) Consecutive failures before leaving as a function of reward number (mean ± s.d.) in example sessions from two different mice. (h) Distribution of the slope coefficients of a linear regression model that predicted the number of consecutive failures before leaving as a function of the number of prior rewards. For visualization, pink are the slope coefficients close to zero (coefficient < 0.5, arbitrary threshold), while blue are sessions with positive slope coefficients. (i) Slope coefficients from (h) between two consecutive sessions (1 and 2) for different mice. Sessions between which the coefficient values vary by more than 0.5 (arbitrary threshold) are highlighted in black. (j) Illustration of the logistic regression model for predicting the switching decision of the mouse from the two different DVs. (k) Deviance explained from the logistic regression that predicts choice behavior based on the DVs (gray box) and from simulated data where the behavior is truly inference-based (white box). On each box, the central mark indicates the median across behavioral sessions, and the bottom and top edges of the box indicate the 25th and 75th percentiles, respectively. The whiskers extend to the most extreme data points. (l) Explained variance from the logistic regression that predicts choice behavior based on the DVs. Sessions where ‘consecutive failures’ is dominant (Var. Exp. Cons. Fail > Var. Exp. Neg. Value) are labeled in pink, while sessions where ‘negative value is dominant’ are labeled in blue (Var. Exp. Cons. Fail < Var. Exp. Neg. Value).

After several days of interaction with this setup (n = 13 ± 5 days; mean ± s.d.), mice (n = 21) learned to exploit each site for several seconds (Fig. 1d,e). Considering the last two sessions of training (n = 42 sessions total), we examined which strategy mice used to time their leaving decisions. As demonstrated previously^8^, for all mice the probability of leaving increased with the number of consecutive failures (Fig. 1f). Yet, not all mice treated rewards equally. For some mice, the number of previous rewards did not affect the probability of leaving after a set number of failures (Fig. 1g, pink), consistent with the inference-based strategy. In contrast, for some other mice the number of failed attempts that they tolerated before leaving the site correlated with the number of previous rewards (Fig. 1g, blue), consistent with the stimulus-bound strategy. We quantified these effects using a linear regression model that predicted the number of consecutive failures before leaving as a function of the number of prior rewards in the current bout (Fig. 1h). We found that the regression coefficient varied strongly within our cohort, consistent with the just-described behavioral heterogeneity across sessions. The distribution across sessions showed signs of bimodality with a dip close to 0.5. Using this criterion, behavior was more consistent with the inference-based strategy in n = 23 sessions (coefficient less than 0.5) and more consistent with the stimulus-bound strategy in the remaining n = 19 sessions (coefficient larger than 0.5). To check if the heterogeneity in strategy was due to variability from session-to-session, mouse-to-mouse, or both, we examined whether the regression coefficients of each mouse varied across consecutive sessions (Fig. 1i). Overall, we observed that the majority of mice kept the same dominant strategy across consecutive sessions (Fig. 1i, gray, but see also Fig. 7), but some mice (n = 4) also switched strategy from one session to the next (Fig. 1i black).

**Figure 7:**
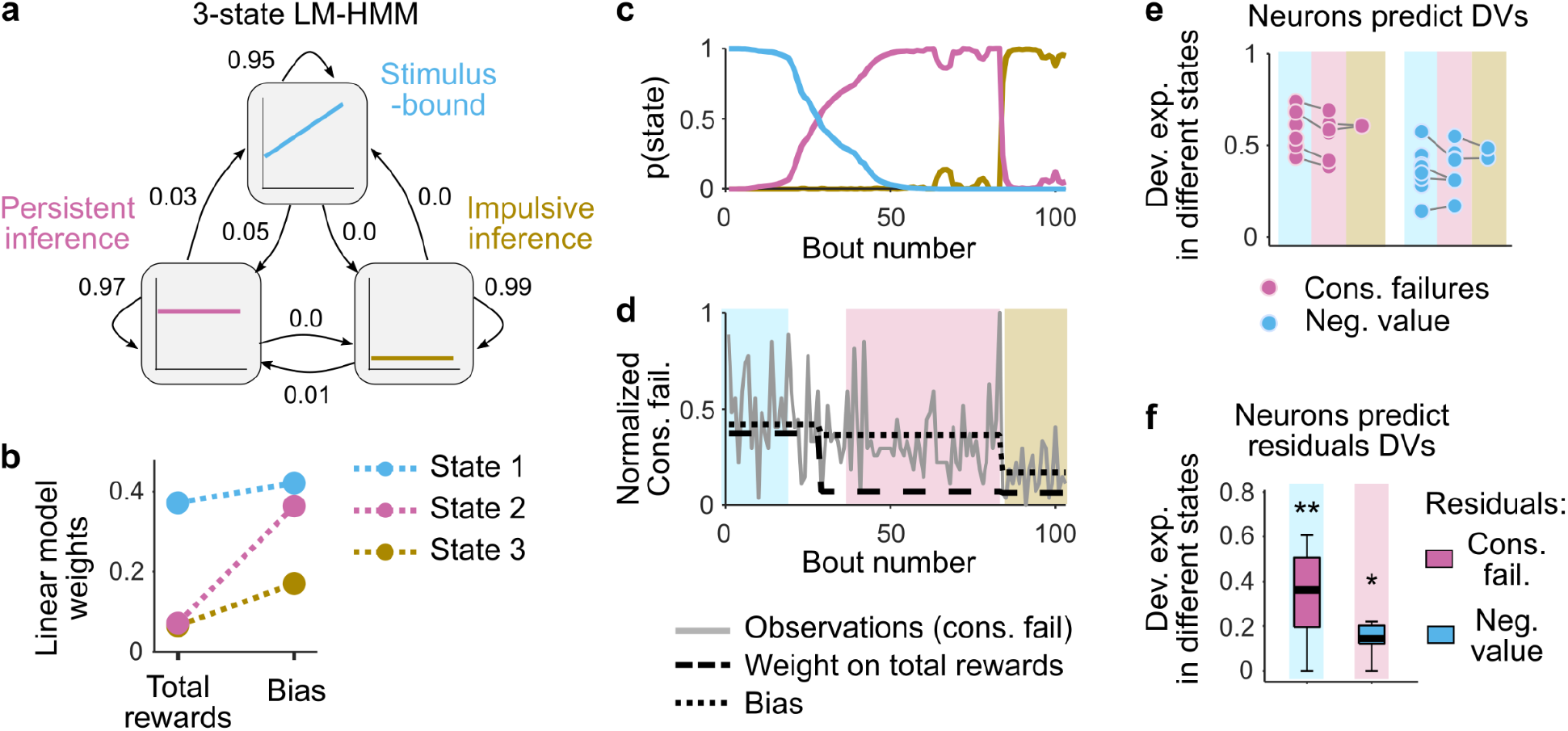
Simultaneous representations of DVs. (a) Illustration of the LM-HMM, with three different states corresponding to different decision-making strategies (labeled ‘Stimulus-bound’, ‘Persistent inference’ and ‘Impulsive inference’). The high self-transition probabilities of 0.95, 097 and 0.99 indicate that states typically persist for many consecutive bouts. The transition probabilities are indicated by the arrows between states. (b) LM weights for the three-state model fit to all sessions simultaneously. (c) Posterior state probabilities (computed with gaussian prior on the weights and Dirichlet prior on transition probabilities) for an example session showing that states typically persisted for many consecutive bouts with high model confidence, but transitioned once or twice over the course of a session. (d) Behavioral data and model parameters of the example session. The gray line indicates the number of consecutive failures (i.e., observations of LMs). The shaded color background indicates the high confidence state (p(state) > 0.75). Dash black lines indicate the LMs bias and weights in each state. (e) Deviance explained from models that fit M2 neurons to the DVs (pink dots: consecutive failures; blue dots: negative value) in different states (high model confidence, indicated by the color background). High confidence intervals were defined as p(state) > 0.75 for at least 25 consecutive bouts. Each dot is a recording session. (f) Deviance explained across sessions (median ± 25th and 75th percentiles) from models that fit M2 neurons to the residual DVs in their respective alternate states of high certainty. Left is the residual consecutive failures (the signal that is orthogonal to the negative value DV) in the stimulus-bound state. Right is the residual negative value (the signal that is orthogonal to the consecutive failure DV) in the inference-based states. Stars indicate that the deviance explained is significantly different from zero (Wilcoxon rank sum test).

These observations indicate that mice vary in their foraging strategies across individuals and sessions, but do not directly indicate how well mice’s behavior are actually described by the DVs. Therefore, we next quantified how well the different DVs ‘consecutive failures’ and ‘negative value’ could predict the precise moment (lick) when an individual mouse would switch sites on a given trial. Specifically, we used regularized logistic regression to model the probability that each lick (n = 2,882 ± 1,631 licks per session; mean ± s.d. across 42 sessions) was the last one in the bout, considering simultaneously the stimulus-bound and the inference-based DVs as predictors (Fig. 1j top, Methods). We estimated the goodness of fit of the two models using the ‘deviance explained’, a generalization of r-squared where ‘0’ meant chance level and ‘1’ meant perfect predictions. We found a median deviance explained of 0.16, a value significantly better than chance for all mice (Fig. 1k, gray box, Wilcoxon rank test: p < 10^-6^). To provide a reference for the meaning of a deviance of this magnitude, we used the same logistic regression model to predict the leaving decisions of a simulated agent in which the ‘ground truth’ was known. For this, we simulated behavioral sessions of an agent making decisions using a logistic function and the DV of the inference strategy with equal numbers of bouts as in the real sessions. We found that the model recovered the ground truth parameters with high accuracy (Extended Data Fig. 2a-d), and performed better than a model attempting to fit the same data using the stimulus-bound DV, which is distinct but correlated with the DV of the inference strategy (Extended data Fig. 2e). Furthermore, the deviance explained by the simulated data (median = 0.25, Extended Data Fig. 2f,g) was only slightly greater than that of the real data (Fig. 1k), indicating that the model with DVs performed close to the maximum that could be expected given the statistical nature of the task. This multivariate approach also confirmed that the two DVs were used to different extents across sessions (Fig. 1l) and, compared to the univariate regression (Fig. 1h), provided even clearer indication of changes in dominant strategy across sessions (Fig. 1l, Extended data Fig. 2h). Finally, the bias term of the model (or intercept) indicated the baseline probability to leave the site (the larger the bias the more impulsive the switching decision), but did not correlate with the use of either DV (Pearson correlation between bias term and explained variance of consecutive failures: *r* = – 0.12, p = 0.44; negative value *r* = – 0.18, p = 0.25).

The logistic regression confirmed that the two DVs describe the switching decision relatively well. Yet, alternative foraging strategies not directly relying on combinations of action outcomes, such as those relying on time since the beginning of the bout or average reward rate, could also explain well the mice’s behavior. Thus, we used the logistic regression model to further explore the space of strategies beyond the two main DVs (Extended Data Fig. 3a). We found that whereas alternative strategies explained some of the behavioral variance, the ‘consecutive failures’ and ‘negative value’ DVs still best predicted the switching decision in most sessions (Extended Data Fig. 3b,c). Although we cannot rule out that mice use other unexplored strategies, these results indicate that the inference-based and stimulus-bound strategies are the best predictors of the switching decision among different classes of foraging strategies.

### Neural activity related to the switching decision

In order to examine the neural basis of DVs underlying the switching decision, we first had to identify brain regions that predicted the switching decision. We used Neuropixels 1.0 electrode arrays^9^, which are single shank probes with hundreds of recording sites that allow registering the activity of large ensembles of neurons (n = 151 ± 59 neurons per session; mean ± s.d.) in multiple regions of the frontal cortex during the task. We targeted the secondary motor cortex (M2; n = 66 ± 37 neurons per session; mean ± s.d.), thought to be important for timing self-initiated actions^10^, planning licking behavior^11^, and predicting changes in behavioral strategy^12^, and the orbitofrontal cortex (OFC; n = 55 ± 24 neurons per session; mean ± s.d.), whose inactivation impacted the performance of inference-based decision-making in freely moving mice in the foraging task^8^. We also recorded in the olfactory cortex (Olf; n = 31 ±23 neurons per session; mean ± s.d.), which is directly ventral to the OFC and can be accessed easily thanks to the long shank of the probe (Fig. 2a,b; Extended Data Fig. 4), but which would not be expected to be specifically involved in this task.

**Figure 2:**
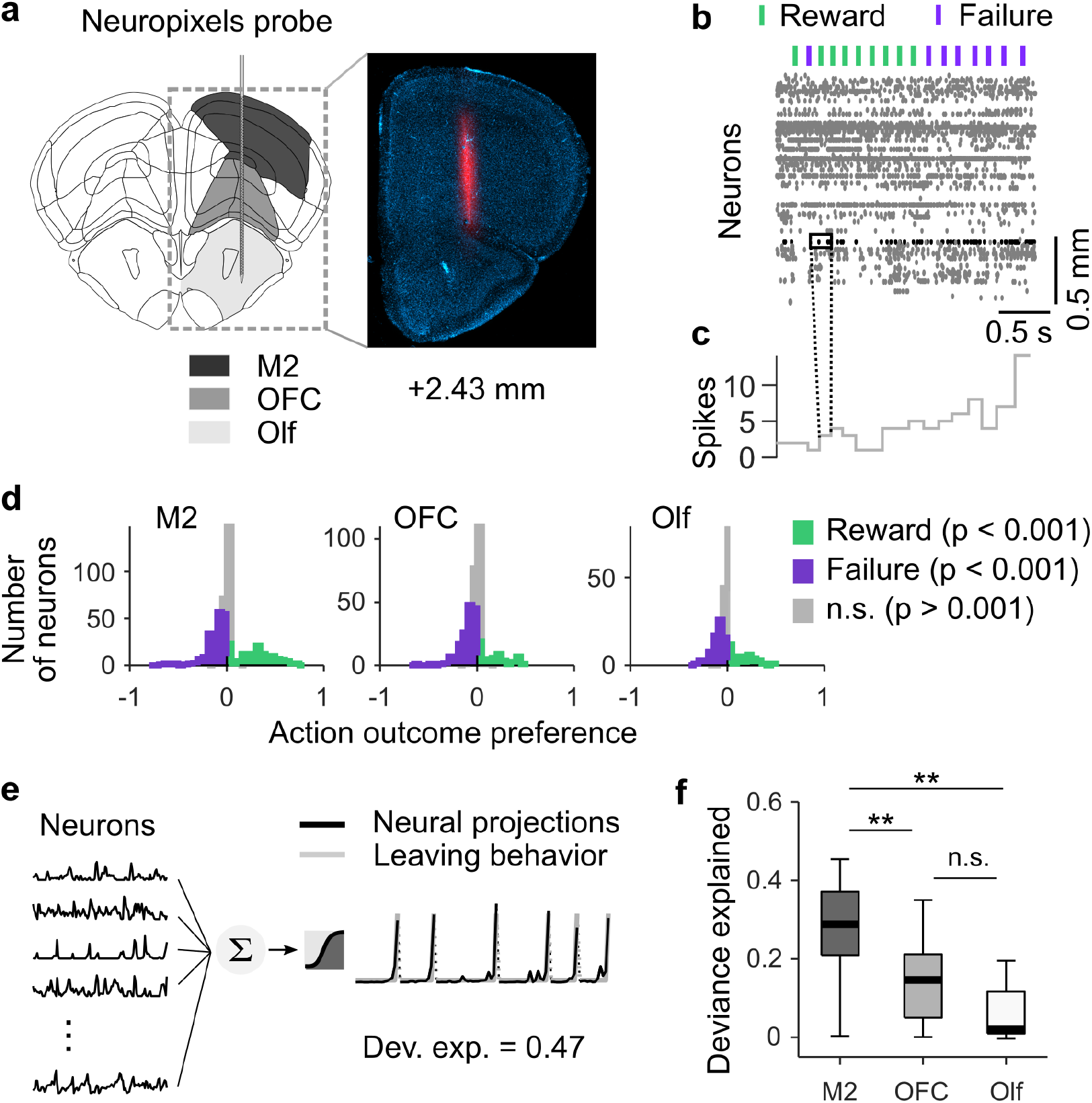
Neural activity related to the switching decision. (a) Schematic target location of probe insertion and an example histology of electrode track. Vertical insertions were performed within a 1 mm diameter craniotomy centered around +2.5 mm anterior and +1.5 mm lateral from Bregma. (b) Example raster plot of 140 simultaneously recorded neurons from M2. Lick-outcome times are indicated by the green (reward) and purple (failure) dashes. (c) Binned response profile of an example neuron. For all analyses, otherwise noted, we averaged for each neuron the number of spikes into bins by considering a 200 ms window centered around each lick. (d) Histogram of outcome selectivity of all neurons recorded M2 (left), OFC (middle) and Olf (right). We used receiver operator characteristic (ROC) analysis to assign a preference index to each neuron. In brief, an ideal observer measures how well the modulation of neuronal firing can classify the outcome (reward or failure) on a lick-by-lick basis. We derived the outcome preference from the area under the ROC curve as: *PREF_R,F_* = 2[*ROC_AREA_*(*f_R_, f_F_*) – 0.5], where *f_R_* and *f_F_* are the firing rate distributions for trials where outcomes are reward and failure respectively. This measure ranges from −1 to 1, where −1 indicates preference for F (failure), 1 means preference for R (reward) and 0 represents no selectivity. The statistical significance of the preference index (p < 0.001, one-sided) was assessed via bootstrapping (1000 iterations). Violet and green bars indicate neurons where the index was significantly different from 0. In all regions, we found neurons significantly modulated by rewards and failures. (e) Illustration of the logistic regression method for predicting the switching decision (gray, right) of the mouse from the principal components of neurons (black, left). (f) Deviance explained from the logistic regression in each region. Two stars indicate a significant difference between regions (Wilcoxon signed rank test: p = 0.0068 between M2 and OFC; p = 0.0049 between M2 and Olf). On each box the central mark indicates the median across recording sessions, and the bottom and top edges of the box indicate the 25th and 75th percentiles, respectively. The whiskers extend to the most extreme data points.

To examine neural responses during the evidence accumulation process, we considered the momentary response patterns of isolated neurons in small time windows (Fig. 2c, Methods). Because we observed heterogeneous task-related activity in many single neurons in all regions (Fig. 2d), we focused on how population activity from each single region predicted the switching decision of mice (n = 11 recording sessions, 1 recording session per mouse except 1 mouse with 2 recording sessions). Using cross-validated and regularized logistic regressions, we decoded the switching decision (i.e., the probability that each lick was the last in the bout) from population responses around each lick (200 ms window) in each session (Fig. 2e, n = 2,533 ± 1,524 licks per session; mean ± s.d. across 11 sessions). To allow for a fair comparison between brain regions, we controlled for the different number of recorded neurons in each region by using as predictors only the first N principal components of neural activity (M2: N = 31 ± 17; OFC: N = 29 ± 9; Olf: N = 16 ± 13), which predicted up to 95% of its total variance (see Methods for additional control analyses). We found that the switching decision could be better decoded using population activity from neurons in M2 than in OFC or Olf (Fig. 2f). This suggests that, unlike OFC, which has been shown to be important for the inference process^8^, M2 may be directly involved in the instantaneous timing of action selection.

### Switching decision and running initiation are dissociable

To test that the neural activity predictive of a switching decision does not simply reflect running initiation, we decoded the switching decision on a subset of behavioral bouts where the last lick and running initiation were clearly decoupled (i.e., they occurred more than 1 s apart; Fig. 3a,b). We found that the last lick could still be decoded with high accuracy, especially in M2 (Fig. 3c), suggesting that M2 activity encodes the intention to switch sites rather than just reflecting the initiation of running behavior.

**Figure 3:**
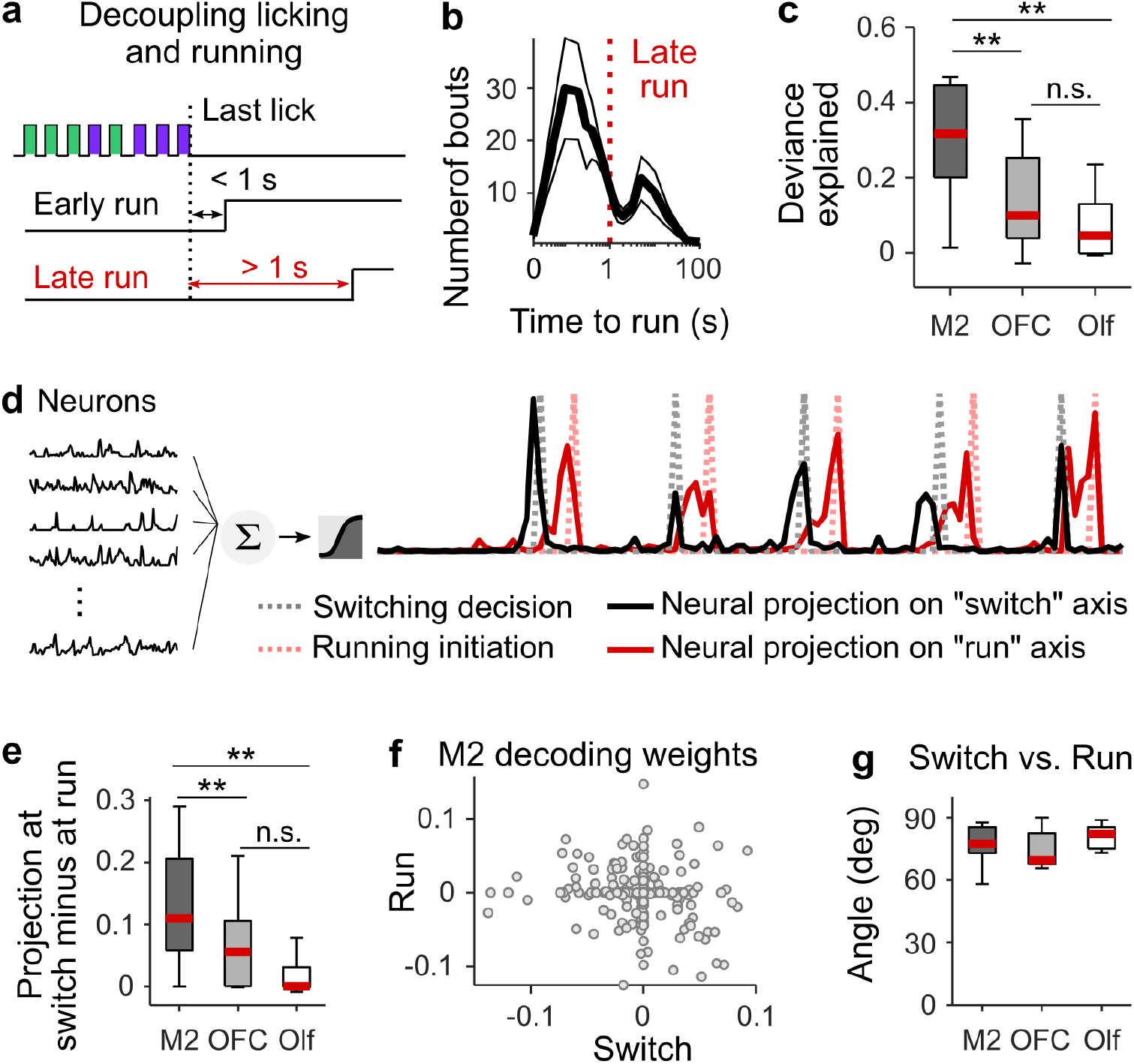
Switching decision and running initiation are dissociable. (a) Last lick always precedes running initiation. Running initiation may occur immediately after the last lick of a bout (< 1 s; ‘early run’) or mice may remain still for several seconds after the last lick and before running initiation (> 1 s; ‘late run’ in red). (b) Bimodal distribution of time between last lick and running initiation (mean ± s.e.m across recording sessions). (c) The deviance explained from models that predict the switching decision (last lick) from the neural activity from M2 (dark gray), OFC (light gray), and Olf (white), in ‘late run’ conditions when the last lick is fully decoupled from running initiation. Two stars indicate a significant difference between regions (Wilcoxon signed rank test: p = 0.002 between M2 and OFC; p = 0.002 between M2 and Olf). (d) Illustration of the logistic regression method for predicting the switching decision (gray dash line) and the running initiation (red dash line) using neural activity from first lick to running initiation (black, left) in bouts when running occurred at least 1 s after the last lick. Red and black solid lines are examples of neural projections onto the two different axes. (e) Difference in values of the neural projection onto the switch axis at the time of switching and the time of running. The larger the difference, the more dissociable the two events. (f) Decoding weights of each M2 neuron (gray dots) for the two different axes. (g) Angles between the two different axes. In all regions, the angle is close to 90 deg indicating that the two axes are closed to orthogonal. On each box of (c), (e) and (g) the central mark indicates the median across recording sessions, and the bottom and top edges of the box indicate the 25th and 75th percentiles, respectively. The whiskers extend to the most extreme data points.

To further test whether the switching decision and running initiation are dissociable in M2, we used neural activity up to the point of running initiation to simultaneously decode the switching decision and the decision to initiate running, again using only bouts where licking and running were decoupled in time (Fig. 3d). The neural activity projected onto the two decoding axes (switching and running) peaked at different times (Fig. 3d,e), and the two axes were close to orthogonal (Fig. 3f,g), consistent with previous studies showing that M2 populations encode preparatory activity for upcoming actions^11,13^. These results indicate that M2 simultaneously encodes, in a separable format, the relevant DVs used to guide an action, as well as a signal associated with the time of initiation of the action itself.

### M2 is involved in the switching decision

The above results point to M2 as a key region for timing the switching decision of the mouse by relying on specific DVs. To further test the contribution of M2 to the implementation of DVs, we partially silenced M2 using an optogenetic strategy (as in Ref^8^). Specifically, we used VGAT-ChR2 mice, which express the excitatory opsin channelrhodopsin-2 in inhibitory GABAergic neurons, to optogenetically silence M2 (especially the superficial layers) bilaterally in 30% of randomly selected bouts (Fig. 4a). We examined 43 sessions from 6 mice, 4 of which were ChR2-expressing and 2 of which were control wild-type littermates implanted and stimulated in the same manner. M2 silencing caused no gross changes in action timing (Extended data Fig. 5), but only a slight decrease in licking rate (Extended data Fig. 5c), and perhaps a trend for the time spent licking (Extended data Fig. 5d). Since M2 inactivation did not significantly impair the motor behavior, we tested if silencing M2 affected the use of the DVs to time the leaving decision. Thus, for each session, we used logistic regressions to estimate how well the DVs predicted the decision to leave the current site during transient inactivation of M2 (Laser ON) and during control bouts (Laser OFF; Fig. 4b). We found that the inactivation of M2 significantly decreased the predictive power of the DVs (Fig. 4c, violet). The same protocol (implantation and laser stimulation) applied to control mice (wildtype littermates that express no inhibitory opsin) had no significant effect on this behavior (Fig. 4c, red). The photostimulation decreased the use of the DV ‘consecutive failures’ (Fig. 4d-f), as well as the leaving bias (Fig. 4g), making animals less inference-based and less impulsive. These results suggest that M2 is part of the neural pathway through which the DVs shape the behavior of the mice.

**Figure 4:**
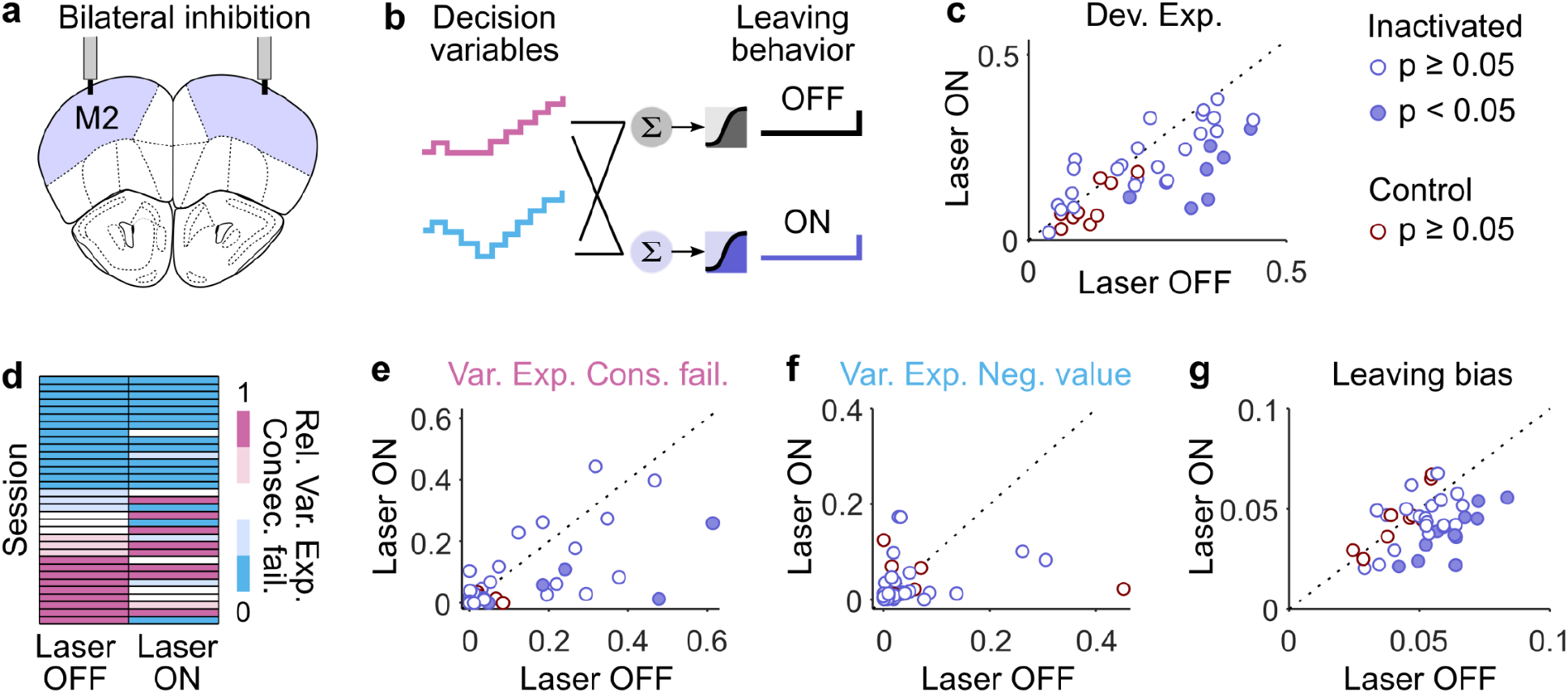
M2 is involved in the switching decision. (a) Schematic target of optic fibers placement. Bilateral photostimulation (5 mW power per fiber, 10 ms pulses at 75 Hz) was triggered by the first lick in 30% of trials and lasted until the last lick of the bout. (b) Illustration of the logistic regression models for independently predicting the switching decision of the mouse based on the DVs during photostimulation (Laser ON) and control bouts (Laser OFF). (c) Deviance explained from the models in (b) for each session (dots) of inactivated mice (violet) and control mice (red) mice. Dots below the identity indicate the sessions where the model performed worse during photostimulation of M2. Filled dots indicate that the effect of photostimulation is significant within single sessions (estimated using resampling, see Methods). To estimate the effect of photostimulation on the deviance explained across mice and session, we used the following mixed model (see methods): Dev.Exp. ~ 1 + Laser + (1 + Laser|Mouse) + (1 + Laser|Session). Fixed effect of stimulation (‘Laser’ predictor): Inactivated: −0.04 ± 0.02, p = 0.021; Control: −0.03 ± 0.014, p = 0.054. (d) Relative variance explained of the DVs for predicting the switching decision in ‘Laser OFF’ vs. ‘Laser ON’ condition. Because both DVs are used as regressors, their relative variances explained sum to 1. Larger values of the relative variance explained of the ‘Consecutive failures’ are colored in pink and indicate that the mouse mainly uses the inference-based strategy. Conversely, lower values of the relative variance explained of ‘Consecutive failures’ are equivalent to larger values of relative variance explained of ‘Negative value’ (colored in blue), indicating the mouse mainly uses the stimulus-bound strategy. (e) Variance explained of ‘Consecutive failures’ in ‘Laser OFF’ vs. ‘Laser ON’ condition. Fixed effect of stimulation: Inactivated: −0.054 ± 0.025, p = 0.032; Control: −0.012 ± 0.009, p = 0.22. (f) Variance explained of ‘Negative value’ in ‘Laser OFF’ vs. ‘Laser ON’ condition. Fixed effect of stimulation: Inactivated: −0.011 ± 0.012, p = 0.35; Control: −0.032 ± 0.045, p = 0.49. (g) Bias term of the logistic regression (intercept) in ‘Laser OFF’ vs. ‘Laser ON’ condition. Fixed effect of stimulation: Inactivated: −0.45 ± 0.078, p < 10^-6^; Control: 0.092 ± 0.075, p = 0.24.

### Neural representation of DVs

The inactivation experiments suggest that one might be able to read out the DV used by the mouse from M2 neural activity, and that M2 might represent this DV better than other cortical regions that afford less accurate predictions of foraging decisions. To test these ideas, we used cross-validated regression-based generalized linear models (GLM; see Methods) to decode the instantaneous magnitude of the DV associated with the behaviorally dominant strategy (i.e. the DV most predictive of behavior) – during each bout (n = 223 ±119 bouts per session; mean ± s.d.) on a lick-by-lick basis – from the responses of neurons (the putative single units) in each recorded brain region (Fig. 5a,b). The example data from Fig. 5a,b, which are from a single recording session during which the dominant strategy of the mouse was the inference (Var. Exp. Con. Fail. = 0.164 vs. Var. Exp. Neg. Value = 0.004), show that the related DV ‘Consecutive failures’ could be decoded with high accuracy from M2 activity (Dev. Exp. = 0.74). In fact, the dominant DV could be well decoded from M2 activity in all sessions (n = 11) from the different mice (Fig. 5c, black). The decodability of dominant DVs was significantly lower in other cortical regions (Fig. 5c; Wilcoxon signed rank test: p < 10^-3^ between M2 and OFC; p < 10^-3^ between M2 and Olf), consistent with the poorer decoding of leaving time in other areas (Fig. 2f).

**Figure 5:**
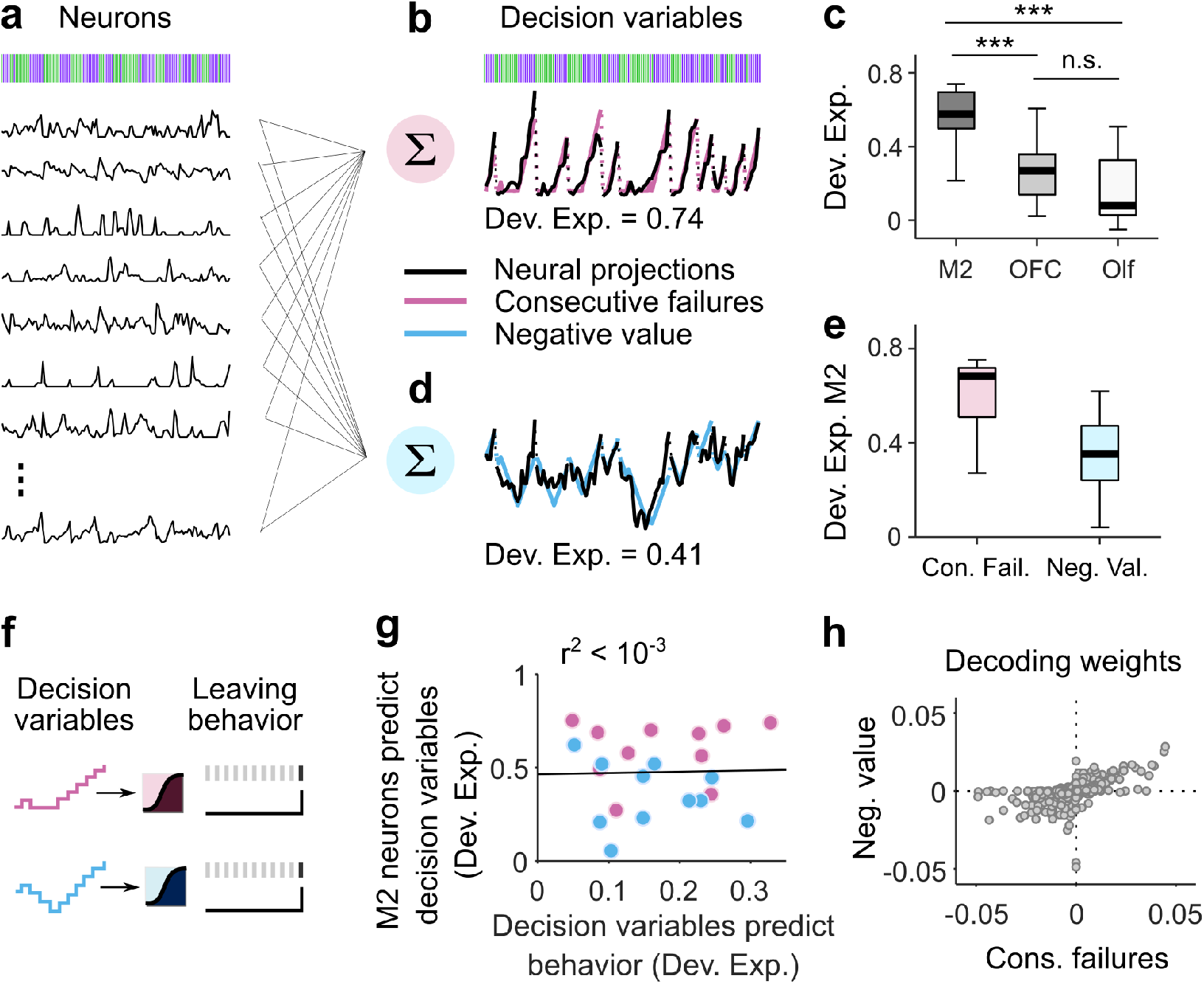
Neural representation of DVs. (a) The regression models take as predictors the activity of simultaneously recorded neurons (black traces) and derive a set of decoding weights to predict the DV. (b) Predictions of the model (black trace is the weighted sums of neural activity) overlaid onto the ‘Consecutive failures’ DV (pink trace). (c) Deviance explained across sessions (median ± 25th and 75th percentiles) from the model in (a,b) in each cortical region. The stars indicate the significance of Wilcoxon signed rank test (p < 0.001). (d) Predictions of the model (black trace is the weighted sums of neural activity) overlaid onto the ‘Negative value’ DV (blue trace). (e) Deviance explained across sessions (median ± 25th and 75th percentiles) predicted from M2 neurons for each DV. (f) Illustration of the logistic regression methods for predicting the switching decision of the mouse from each DV separately. (g) Correlation between the neural representations of different DVs (color coded as in b,d) and how well each DV predicts behavior. Each dot corresponds to a particular DV from a given recording session. The linear regression is reported in black. (h) Decoding weights of each M2 neuron (gray dots; total across recording n = 778) for the two different DVs.

Since we have shown that different mice can rely on different DVs and individual mice can change decision strategies across sessions (Fig. 1), we next asked whether session-by-session heterogeneity in decision strategy could be explained by the degree to which M2 neurons reflected the DVs in a given session. Here, we used the GLM to compare the decoding of the dominant and the alternative DVs from M2 neurons in each recording session (Fig. 5a,d). Contrary to our expectation, we found that decoding was actually similar between the dominant and alternative decision-strategies. For instance, in the example session of Fig. 5a,b,d, despite the selectivity of the behavior for inference-based decisions, the DV supporting the stimulus-bound strategy could also be well decoded from M2 (Dev. Exp. = 0.41). This finding was consistent across our experiments: in all sessions, the DVs could both be read out from M2 activity (Fig. 5e; Extended Data Fig. 6). On average, the ‘Consecutive failures’ DV was somewhat better represented than the ‘Negative value’ (Wilcoxon signed rank test: p < 10^-3^). This average difference could stem from the fact that the majority of mice (8 out of 11) used the inference-based strategy that relies on the ‘Consecutive failures’. Thus, to test whether the DV that was most predictive of the switching decision was also the one that was better decoded from M2 on a session-by-session basis, we predicted the decision to switch sites from each DV (Fig. 4f) and compared the accuracy of this prediction to the accuracy of the neural representations of the DVs (Fig. 5g). There was no correlation between how M2 represented each DV in a session and how well the DV predicted behavior in the same session (r^2^ < 10^-3^, p = 0.9). Therefore, together these analyses suggest that whereas M2 neural activity is important to the execution of a decision strategy (Fig. 4), the pattern of neural activity in M2 is not adapted to represent specifically the DV executed by the mouse, and instead reflects a broader range of decision strategies even when they are not currently used.

To further characterize the multiplexing of DVs in M2, we asked whether different variables are supported by distinct or overlapping populations. In particular, we examined the weights assigned to each neuron when decoding the ‘Consecutive failures’ vs. the ‘Negative value’ (Fig. 5h). We found that decoding weights for both DVs were strongly correlated (Pearson coefficient = 0.56, p < 10^-4^), indicating a considerable overlap between the populations of M2 neurons that supported each DV, as opposed to compartmentalization into distinct populations for each variable.

### Independent representations of DVs

A possible concern with the interpretation that M2 multiplexes actual (used) and ‘counterfactual’ (unused) DVs is that alternative DVs might be decodable only by virtue of being similar to the one reflected behaviorally. Although the computations underlying the two DVs are different, for the particular sequences of rewards and failures experienced by the mice, the DVs themselves are indeed fairly correlated overall (Pearson coefficient: 0.79 ± 0.15; mean ± s.d.).

As a first strategy to overcome this limitation, we took advantage of the previously noted fact that the two different DVs being considered are the same in the way they treat failures but differ in the way that they treat rewards: while the ‘negative value’ requires negative integration of rewards, the ‘consecutive failures’ variable requires a complete reset by a single reward (Fig. 6a). Analysis of subsets of sequences that consist of multiple consecutive rewards should therefore reveal the differences between the two DVs (Fig. 6b). To test this, we sub-selected lick sequences and sorted them according to the relative number of rewards and failures. This produced subsequences with varying degrees of correlation between the two decision-variables (Fig. 6c). We then ran the same decoding analyses as before on these subsequences of M2 activity. We found that the ability to decode the subsequences was actually independent of their degree of correlation (Fig. 6d, one-way ANOVA for each sequence across correlation values followed by multiple pairwise comparison tests, all p-values > 0.05). Our second approach was to investigate whether we could specifically decode the component of each DV that is uncorrelated with the other one, i.e., its residual. Indeed, we could decode the residuals from both DVs from the activity of M2 populations (Fig. 6e-f). Together, these results establish that the ability to decode an alternative DV does not arise from the correlation from that variable with the dominant DV. Interestingly, this approach revealed that OFC better represented the ‘Consecutive failures’, consistent with previous work suggesting that OFC is important for the inference-based strategy.^8^

**Figure 6:**
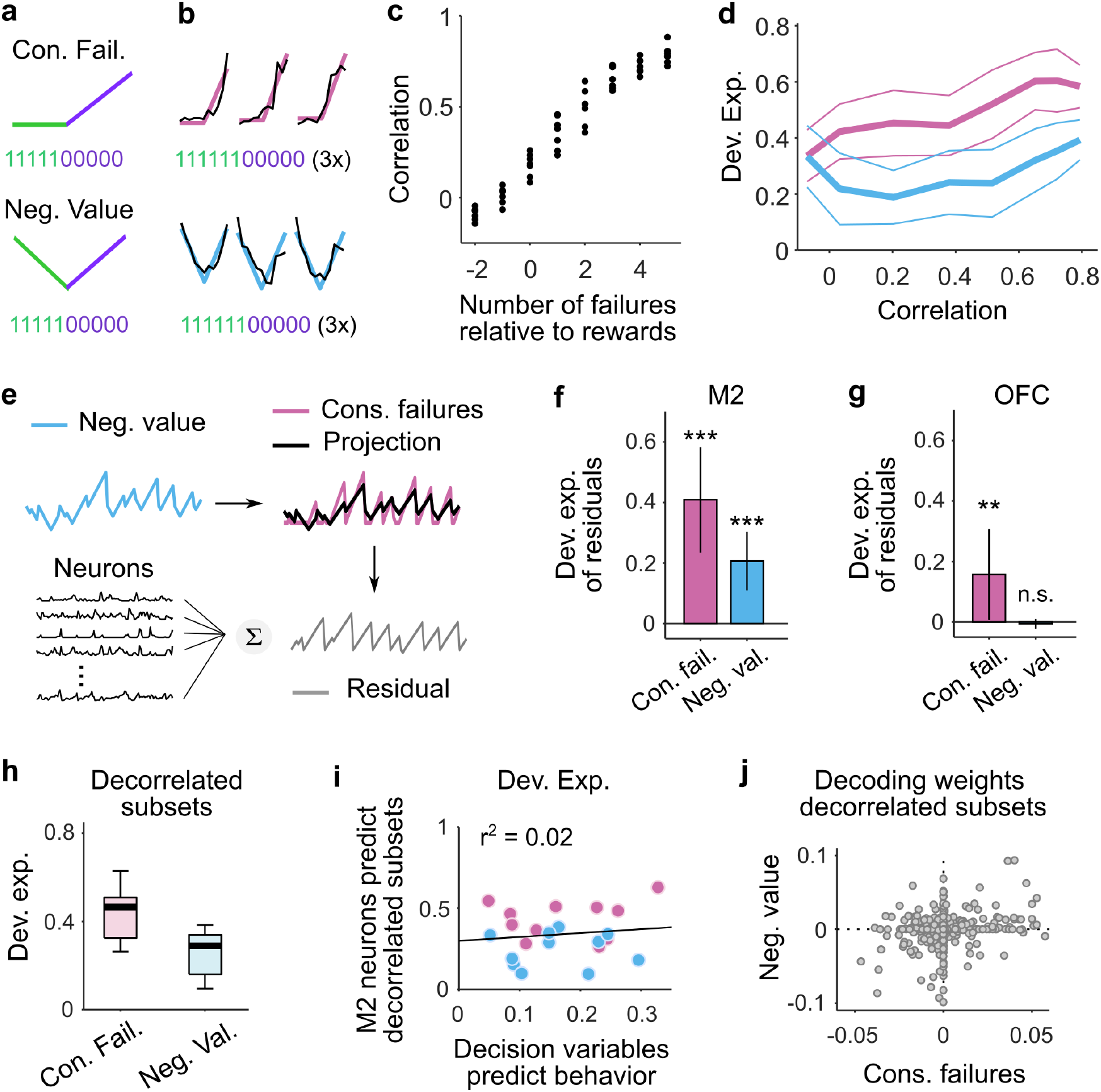
Independent representations of DVs. (a) Two different sequences relying on different computations involving reset (top) and accumulations (bottom) of rewards. (b) Three example bouts (columns) of population activity (black traces) projected onto the dimensions that best predict the trajectory of the different sequences (color traces). Only subsequences of consecutive rewards followed by consecutive failures were selected in order to visualize the different computations in (a) (~5% of bouts). (c) Selecting subsets of action outcomes where the total number of failures changes relative to the number of rewards (abscissa) alter the correlation between sequences generated with the computations in (a) (ordinates). Black dots for each value of the number of failures represents a recording session. (d) How well the sequences relying on the two different computations can be decoded from M2 (ordinates) as a function of the correlation between them (median ± MAD across sessions). Pink are sequences that accumulate failures and reset with rewards (equivalent to ‘consecutive failures’). Blue are sequences that accumulate failures upward and rewards downward (equivalent to ‘negative value’). (e) Schematic description of our strategy to linearly regress each of the two DVs on the other. This approach allowed us to express DV1 (e.g., ‘Consecutive failures’) as the sum of a time-series proportional to DV2 (e.g., ‘Negative value’) plus a time-series orthogonal (uncorrelated) to DV2, which we denote as its residual. Here, the ‘consecutive failures residual’ (gray) is orthogonal to the ‘negative value’ (blue). The same procedure was used to generate the ‘negative value residual’ orthogonal to the ‘consecutive failures’. Both residuals were then fit by M2 neurons. (f) Deviance explained across sessions (median ± mad) of the model in (e). Pink: residual consecutive failures; Blue: residual negative value. The residuals relative to each decision variable were both significantly represented in M2 (Wilcoxon rank sum test: p < 10^-3^ for both, indicated by the stars). The size of the pink bar measures how well one can decode the part of ‘Consecutive failure’ orthogonal to ‘Negative value’ (residual consecutive failures) and the size of the blue bar measures how well one can decode the part of ‘Negative value’ orthogonal to ‘Consecutive failure (residual negative value)’. If only ‘Consecutive failure’ were represented, the residual consecutive failures should be represented, but the residual negative value would not be represented. On the other hand, if both DVs are represented, both residuals should be decodable, as shown here in M2. (g) Same as in (f) but with OFC neurons. The residuals ‘Consecutive failures’ were decodable from OFC ensembles (pink; Wilcoxon rank sum test: p = 0. 0029), but the residuals ‘Negative value’ were not (pink; Wilcoxon rank sum test: p = 0. 52). (h) Deviance explained across sessions (median ± 25th and 75th percentiles) predicted from M2 neurons for each decorrelated subsets of DVs (Wilcoxon signed rank test: p < 10^-3^). (i) Correlation between the neural representations of decorrelated subsets of DVs (color coded as in b,d,e) and how well each DV predicts behavior. Each dot corresponds to a particular DV subsets from a given recording session. The linear regression is reported in black (r^2^ = 0.02, p = 0.6). (j) Decoding weights of each M2 neuron (gray dots; total across recording n = 778) for the decorrelated different subset of DVs (Pearson coefficient between decoding weights = 0.20, p < 10^-7^).

Using only the particular sequences of trials in which the DVs were fully decorrelated (Pearson correlation between DVs: 0.03 ± 0.02; median ± MAD across session), we again tested the possibility that the DVs that were best decoded from M2 were the most predictive of behavior (as in Fig. 5e,g,h). Here, the ‘Consecutive failures’ remained better represented than the ‘Negative value’ (Fig. 6h). Similar to the results with the intact DVs, there was no correlation between how M2 represented each decorrelated subset of DVs and how well the DV predicted behavior (Fig. 6i). This was the case even if the populations of M2 neurons that supported each decorrelated subset of DVs were nearly orthogonal, as indicated by the small correlation between decoding weights (Fig. 6j).

### DV multiplexing does not reflect fast switching between strategies

While one interpretation of multiplexing is true simultaneous representation of multiple DVs, our interpretation is relying on decoding analyses carried out over entire sessions of behavior. Could it be that multiplexing of DVs actually results from sequential switching between the two strategies within a session? To investigate this, we first examined whether there was any evidence that mice switched strategies within a session using a framework based on hidden Markov models (HMM) combined with linear regression models (LM) (see methods and Ref^14^). The resulting ‘LM-HMM’ framework modeled the number of consecutive failures that the animal bears before switching sites using two inputs: (1) the total number of rewards, which allows distinguishing between inference-based (i.e. reward independent) and stimulus-bound (i.e. reward dependent) strategies, as in Fig. 1g, and (2) a constant bias, which reflects the level of impulsivity of the animal. Each hidden state in the model captures a specific dependence of consecutive failures on the total rewards and the bias, characterizing a particular decision-making strategy.

A model with three states best described the switching decision and yielded interpretable and persistent states (Fig. 7a, Extended Data Fig. 7a). One of the states had large weight on the number of rewards, indicative of a stimulus-bound strategy, while the other two had negligible weights on rewards, consistent with the inference (Fig. 7b, Extended Data Fig. 7b,c). To visualize the temporal structure of the foraging decision within a session, we computed the posterior probability over the latent states across all behavioral bouts (Fig. 7c,d), which revealed that mice mostly remained in discrete states (average probability of the dominant strategies over all bouts: 0.91 ± 0.06; median ± MAD across sessions) for many bouts in a row (average duration of states: 56 ± 41 bouts; median ± MAD across sessions), but tended to switch states at least once per session (state transition in 8 sessions out of 11; Extended Data Fig. 7d).

Since mice alternated between states of inference-based and stimulus-bound strategies within the course of their recording session, we examined whether we could decode better from M2 activity the ‘consecutive failures’ DV during the inference-based states than during the stimulus-bound states (Fig. 7e, pink dots), and vice versa for ‘negative value’ DV (Fig. 7e, blue dots). Consistent with the whole-session analysis (Fig. 5g), there were no significant differences between how well a given DV could be decoded when the mice’s behavior relied on it or when it did not (Wilcoxon rank sum test: p > 0.05 for all comparisons). The residual signals after the DVs, which are orthogonalized, were also decodable in their respective alternate states (Fig. 7f). These analyses suggest that multiplexing of strategy is not due to switch of strategies within a session.

### M2 represents foraging algorithms

Given that M2 appears to multiplex different DVs, we wondered whether this might reflect a generic capacity to represent any signal with similar temporal characteristics as the DVs in the task, as predicted by the reservoir computing framework^15–17^. Decoding analyses of artificial signals with matched temporal statistics revealed this not to be the case (Extended Data Fig. 8). Therefore, we next considered that the space of signals encoded in M2 might be restricted to potentially meaningful variables generated from a common set of essential computations. Here, the two DVs we have been considering could both be conceptualized as an adaptive, outcome-dependent feedback gain on a running count. For instance, if we refer to the running count after the *t-th* lick as *x_t_* and to the outcome of the next lick as *o*_*t*+1_ (equal 1 or 0 if the outcome is a reward or a failure respectively), then we can write the update rule compactly as

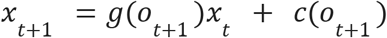

with *g*(*o*_*t*+1_ = 1) =0, *g*(*o*_*t*+1_ = 0) = 1, and *c*(*o*_*t*+1_ = 1) = *c*(*o*_*t*+1_ = 0) = 1 for the inference-based DV, and *g*(*o*_*t*+1_ = 1) = *g*(*o*_*t*+1_ = 0) = 1, and *c*(*o*_*t*+1_ = 0) = – *c*(*o*_*t*+1_ = 1) = 1, for the stimulus-bound DV. This realization suggests that a common generative model, which we named the ‘INTEGRATE-AND-RESET model’, can produce these two different DVs by adjusting certain model parameters (Fig. 8a). The INTEGRATE-AND-RESET model describes, within a single algorithmic framework, the computations necessary to generate, not only the two DVs considered so far, but also other DVs relevant for a variety of other commonly studied behavioral tasks. For instance a ‘global count’ (accumulated number of outcomes) DV is related to counting or timing tasks^18,19^. Similarly, matching tasks involving randomly timed, cached rewards, are optimally solved by integrating the difference between rewards and failures with an exponential decay^20^. Sequential foraging in patchy environments is also solved by integrating the difference between rewards and failures, equivalent to tracking the relative ‘negative value’ of a foraging site^21^. Other integration tasks, like the ‘poisson clicks’ task^22^, require perfect integration of two variables. Thus, the space of DVs generated by the INTEGRATE-AND-RESET model covers a large space of tasks that have been studied in the lab and might be useful in different behavioral contexts.

**Figure 8:**
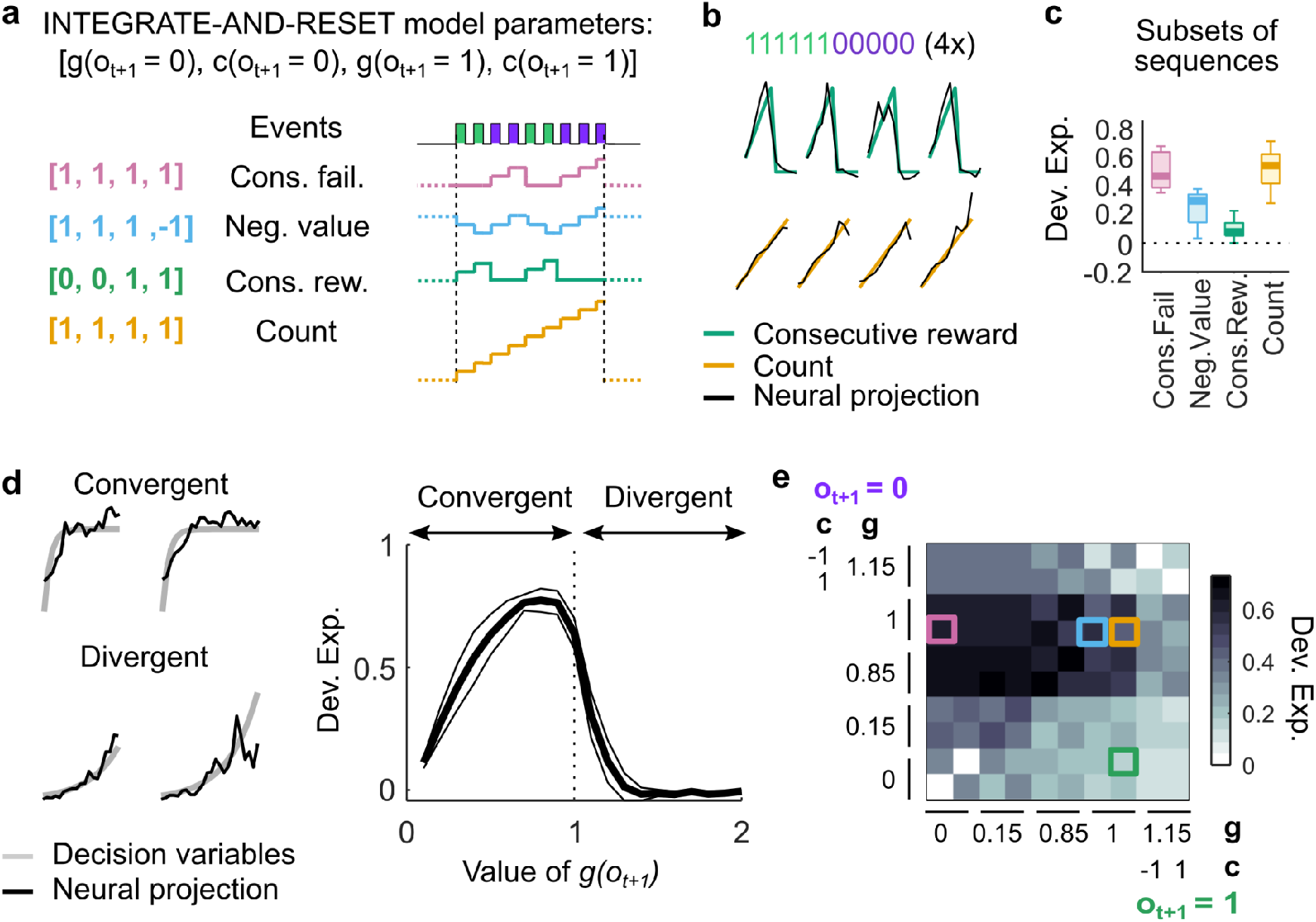
M2 represents foraging algorithms. (a) The INTEGRATE-AND-RESET model with four parameters generates different time series by accumulating, resetting or ignoring each possible event (reward or failure). In the simplest instantiation of this model, the two outcome-dependent parameters are discrete: one is a gain factor (*g*) that specifies whether the running count should be reset or accumulated by each outcome – a non-linear operation – and the other (*c*) specifies how each outcome linearly contributes to the resulting running count, which in general could be positive, negative, or zero (leaving it unaffected; see Methods for more details). Each specification of these two discrete parameters leads to a different DV Example set of parameters yielding example DVs on the right. (b) Four example bouts (columns) of population activity (black traces) projected onto the dimensions that best predict the trajectory of the ‘Consecutive rewards’ (green) and ‘Count’ (yellow). Only subsequences of consecutive rewards followed by consecutive failures were selected to highlight the computations underlying the different variables. (c) Deviance explained across sessions (median ± 25th and 75th percentiles) of the 4 basis sequences decoded from M2 population activity. The sequences were decorrelated using the same method as in Fig. 6c,d. (d) Left: example sequences (gray) produced by analog parameters (Convergent: c(o_t+1_ = 0) = c(o_t+1_ = 1) = 1 and g(o_t+1_ = 0) = g(o_t+1_ = 1) = 0.5; Divergent: c(o_t+1_ = 0) = c(o_t+1_ = 1) = 1 and g(o_t+1_ = 0) = g(o_t+1_ = 1) = 1.15). Black traces are the neural projection from M2 population activity. Right: deviance explained from decoding sublinear and supralinear integrations by M2 population activity. Here, we show an example where the parameters of the INTEGRATE-AND-RESET model are: c(o_t+1_ = 0) = c(o_t+1_ = 1) = 1 and g(o_t+1_ = 0) = g(o_t+1_ = 1). (e) Matrix of deviance explained from decoding sequences with different time constants (corresponding to different values of *g*) of integrations of rewards (columns) and failures (rows) with M2 population activity. The basis sequences are indicated by the color-coded squares.

All non-trivial time series produced by the INTEGRATE-AND-RESET model can be expressed as linear combinations of four basis sequences (Fig. 8a; Methods). The two sequences involving reset describe integration of failures and reset by rewards (‘consecutive failures’) and vice-versa (‘consecutive rewards’). The two sequences for accumulation without reset are upwards integration of both rewards and failures (equivalent to ‘count’) and integration upwards of rewards and downwards of failures (equivalent to ‘negative value’). We already know that M2 simultaneously represents two of these basis elements (‘consecutive failures’ and ‘negative value’). Thus, we tested whether M2 also represented the two additional basis sequences. We found that, indeed, ‘Consecutive reward’ and ‘Count’ could be decoded from the M2 population (Fig. 8b, Wilcoxon signed rank test: p < 10^-3^ for both), and remained decodable from the M2 population when using the subsequences that decorrelate the variables (Fig. 8c; Wilcoxon signed rank test: p = 0.002 for ‘Consecutive reward’ and p < 10^-3^ for ‘Count’). In particular, the sequences that consisted of integration of both reward and failures in the same direction (‘Count’) could be recovered with high accuracy.

The INTEGRATE-AND-RESET model can be extended, through analog values of ‘g’, to produce sequences with different dynamics and various time constants (Fig. 8d left). Note that adjusting analog parameter values can directly relate the INTEGRATE-AND-RESET model to frameworks of reinforcement learning with differential learning, where the “reset” is equivalent to a very large negative rate of decay. Therefore, we further tested the richness of the actual INTEGRATE-AND-RESET model family instantiated by M2 by decoding sequences generated with analog ‘g’. We found that M2 could also represent leaky integration of rewards and failures, and even amplification with small positive feedback (g(o_t+1_) < 1.2, Fig. 8d, right). Comparing across this parameter space (Fig. 8e), we observed that M2 had a preferred mode of integration that consisted of mostly perfect integration of failures (0.85 ≤ g(o_t+1_ = 0) ≤ 1) and integration of rewards with a variety of time constants (g(o_t+1_ = 1) ≤ 1). Altogether, our results show that M2 simultaneously represents a relatively large repertoire of computations that embody a variety of foraging DVs, potentially spanning a set of optimal strategies for environments with different dynamics for the latent state.

## DISCUSSION

We explored the capacity of several regions of the cortex to deploy different algorithms for generating a diversity of DVs. We studied this in the context of a foraging task whose solution required mice to process streams of successful and unsuccessful foraging attempts (licks) executed over several seconds. We found that mice could use not one but a set of discrete processing strategies to time their decision to switch between foraging sites, and our newly developed LM-HMM framework revealed that mice often change strategies within a session. All of the decision strategies could be well read out from populations of neurons in M2. Moreover, we found the set of potentially relevant DVs were actually implemented in parallel within the same neural populations in M2. Conversely, OFC did not appear to multiplex DVs, consistent with the idea that it may be specifically involved in the computations of the inference-based strategy^8^.

While ‘causal’ manipulations of M2 using optogenetic inactivation showed that M2 was important to the deployment of the inference-based strategy, we found that the neural availability of alternative DVs was nearly independent of the actual behaviorally-deployed DV. Functionally, the ability of M2 to multiplex the computation of several DVs could allow the mice to rapidly explore and adapt behavior to dynamically changing environmental contingencies by simply modifying linear readouts of M2 neural populations^23,24^ without the need to implement new computations.

The different DVs in M2 were “mixed” but could be recovered through linear decoding. Although multiplexed neural codes have been observed previously in other cortical regions^15,25–28^, our results establish that the kind of information that is multiplexed is not limited to representations of instantaneously observable events in premotor regions, but also includes temporally extended computations spanning several seconds. While the observation of multiplexed DVs is reminiscent of the framework of ‘reservoir’ computing^15–17,29^, we found that M2’s coding capacity was not universal, and instead specifically implemented a substantial but specifically circumscribed pool of potentially meaningful computations. One computation is accumulation of evidence, which, through its intimate relationship with posterior beliefs^30,31^, constitutes an essential computation for statistical inference, and has therefore been implicated in a variety of decision-making and reasoning tasks^22,32–36^. Accumulation (possibly temporally discounted) of action outcomes also underlies several reinforcement learning algorithms^37–40^. Although less attention has been devoted to reset-like computations (but see Ref^41^), they are also essential for inference when certain observations specify a state unambiguously^8^.

The two strategies that we describe in the context of foraging represent a particular example of a more general phenomenon. In complex environments, agents can adapt their behavior in different ways depending on how accurately they can infer and specify the relevant causal structure^42^, a process which, in the lab, can be described as finding the correct “task representation”. Even if unable to apprehend the true causal model, agents can display reasonably well adapted behavior by leveraging the predictive power of salient environmental events. However, because the task representation is not correct, the association between these events and outcomes will necessarily be more probabilistic from the point of view of the agent. Such agents incorrectly model unexplained outcome variance (arising from incomplete task representations) as unexplainable, and often resort to exploratory strategies that are adaptive in what they construe as highly volatile environments^43,44^. Our stimulus-bound strategy is an example of this phenomenon. Lapsing in psychophysical discrimination tasks – i.e, errors that cannot be attributed to sensory limitations – can also be interpreted as exploratory choices that are made when a failure to aprehend the true stimulus dimension that needs to be discriminated is interpreted by the agent as evidence for probabilistic action-outcome mappings^45^. Our results suggest that, at least in the case of foraging, the computations necessary to implement strategies lying along this continuum are computed simultaneously and available, which might facilitate the process of “insight” necessary to switch between them.

Our finding also speaks to the debate on the nature of serial processing limitations in the brain. While it has been shown that limitations apply in some kinds of evidence accumulation tasks^2,4,46^, here we show in a different, but ethologically important, setting that some forms of evidence accumulation can proceed in parallel. An important difference between our task and standard behavioral paradigms that study cognitive bottlenecks is that our mice do not need to simultaneously compute two DVs to perform the task successfully. Nevertheless, we show unambiguously that neural populations in the premotor cortex of mice using a strategy where a single reward resets a counter of failures, reveal both this reset, and simultaneously the updating of a reward counter. Our findings are thus consistent with proposals favoring parallel integration^47,48^ and with models that place serial constraints on behavior close to the specification of the timing of action^47,49^.

## METHODS

### Animal subjects

A total of 30 adult male and female mice (24 C57BL/6J and 6 VGAT, 2-9 months old) were used in this study. All experimental procedures were approved and performed in accordance with the Champalimaud Centre for the Unknown Ethics Committee guidelines and by the Portuguese Veterinary General Board (Direco-Geral de Veterinria, approval 0421/000/000/2016). During training and recording, mice were water-restricted (starting 5 to 10 days after head-bar implantation), and sucrose water (10%) was available to them only during the task. Mice were given 1 mL of water or 1 gram of hydrogel (Clear H2O) on days when no training or recording occured or if they did not receive enough water during the task.

### Surgery and head-fixation

All surgeries used standard aseptic procedures. Mice were deeply anesthetized with 4% isoflurane (by volume in O_2_) and mounted in a stereotaxic apparatus (Kopf Instruments). Mice were kept on a heating pad and their eyes were covered with eye ointment (Vitaminoftalmina A). During the surgery, the anesthesia levels were adjusted between 1% and 2% to achieve 1/second breathing rate. The scalp was shaved and disinfected with 70% ethanol and Betadine. Carprofen (non-steroidal anti-inflammatory and analgesic drug, 5 mg/kg) was injected subcutaneously. A flap of skin (less than 1cm^2^) was removed from the dorsal skull with a single cut and the skull was cleaned and dried with sterile cotton swabs. The bone was scraped with a delicate bone scraper tool and covered with a thin layer of cement (C&B super-bond). Four small craniotomies were drilled (HM1 005 Meisinger tungsten) between Bregma and Lamba (around −0.5 and −1 AP; ±1 ML) and four small screws (Antrin miniature specialities, 000-120×1/16) previously soaked in 90% ethanol, were inserted in the craniotomies in order to stabilize the implant. The head-bar (stainless steel, 19.1 × 3.2 mm), previously soaked in 90% ethanol, was positioned directly on top of the screws. Dental cement (Tab 2000 Kerr) was added to fix the head-bar in position and to form a well around the frontal bone (from the head-bar to the coronal suture). Finally an external ground for electrophysiological recording (a male pin whose one extremity touched the skull) was cemented onto the head-bar.

### Behavioral apparatus

Head-fixed mice were placed on a linear treadmill with a 3D printed plastic base and a conveyor belt made of Lego small tread links. The running speed on the treadmill was monitored with a microcontroller (Arduino Mega 2560), which acquired the trace of an analog rotary encoder (MAE3 Absolute Magnetic Kit Encoder) embedded in the treadmill. The treadmill could activate two movable arms via a coupling with two motors (Digital Servo motor Hitec HS-5625-MG). A lick-port, made of a cut and polished 18G needle, was glued at the extremity of each arm. Water flowed to the lick-port by gravity through water tubing and was controlled by calibrated solenoid valves (Lee Company). Licks were detected in real time with a camera (Sony PlayStation 3 Eye Camera or FLIR Chameleon-USB3) located on the side of the treadmill. Using BONSAI^50^, an open-source visual programming language, a small squared region of interest was defined around the tongue. To detect the licks a threshold was applied to the signal within the region of interest. The behavioral apparatus was controlled by microcontrollers (Arduino Mega 2560) and scientific boards (Champalimaud Hardware platform), which were responsible for recording the time of the licks and the running speed on the treadmill, and for controlling water-reward delivery and reward-depletion according to the statistics of the task.

### Task design

In the foraging task, two reward sites, materialized by two movable arms, could be exploited. Mice licked at a given site to obtain liquid reward and decided when to leave the current site to explore the other one. Each site could be in one of two states: “ACTIVE”, i.e. delivering probabilistic reward, or “INACTIVE”, i.e. not delivering any reward. If one of the sites was “ACTIVE”, the other one was automatically “INACTIVE”. Each lick at the site in the “ACTIVE” state yielded reward with a probability of 90%, and could cause the state to transition to “INACTIVE” with a probability of 30%. Licks could trigger the state of the exploited site to transition from “ACTIVE” to “INACTIVE”, but never the other way around. Importantly, this transition was hidden to the animal. Therefore, mice had to infer the hidden state of the exploited site from the history of rewarded and unrewarded licks (i.e., reward and failures). We defined ‘behavioral bout’ the sequence of consecutive licks at one spout. A tone (150 ms, 10 kHz) was played when one of the arms moved into place (i.e., in front of the mouse) to signal that a bout could start. At the tone, the closed-loop between the motors and the treadmill decoupled during 1.5 s or until the first valid lick was detected. During this time mice had to “STOP”, i.e. decrease their running speed for more than 250 ms below a threshold for movement (6 cm/s). Licks were considered invalid if they happened before “STOP” or at any moment after “STOP” if the speed was above the threshold. If a mouse failed to “STOP”, “LEAVE” was triggered by reactivating the closed-loop after 1.5 s, which activated the movement of the arms (the one in front moved away and the other moved into place). Mice typically took around 200 ms to “STOP” and initiate valid licking. During the licking periods, each lick was rewarded in a probabilistic fashion by a small drop of water (1 μl). The small reward size ensured that there was no strong difference in licking rate between rewarded and unrewarded licks. To “LEAVE”, mice had to restart running above the threshold for movement for more than 150 ms, and travel a fixed distance on the treadmill (around 16 cm) to reach the other arm. We defined as correct bouts the ones in which mice stopped licking after the states transitioned from “ACTIVE” to “INACTIVE”. Error bouts were ones in which mice stopped licking before the state transition occurred. In this case, mice had to travel double the distance to get back to the arm in “ACTIVE” state. Missed bouts were ones in which mice alternated between arms without any valid lick. These ‘missed bouts’ were excluded from our analysis.

### Mouse training

Mice were handled by the experimenter from 3 to 7 days, starting from the beginning of the water restriction and prior to the first training session. At the beginning of the training, mice were acclimatized to the head-fixation and to the arm movement and received liquid reward simply by licking at the lick-port. The position of the lick-ports relative to the snout of the mouse had an important effect on behavioral performances. Thus, to ensure that the position of the lick-ports remained unchanged across experimental sessions, it was carefully adjusted on the first session and calibrated before the beginning of every other session. There were no explicit cues that allow discriminating between the two arms, and it was not even necessary that animal be fully aware of the two different arms to perform the task. After mice learned to lick for water reward (typically after one or two sessions), the next sessions consisted of an easier version of the task (with low probability of state transition, typically 5% or 10%, and high probability of reward delivery, 90%), and both arms in “ACTIVE” state. That way, if mice alternated between arms before the states of the sites transitioned, the other arm would still deliver reward and animals would not receive the travel penalty. Occasionally during the early phase of training, manual water delivery was necessary to motivate the mice to lick or to stop running. Alternatively, it was sometimes necessary to gently touch the tail of the animals, such that they started to run and gradually associated running with the movement of the arms. The difficulty of the following sessions was progressively increased by increasing the probability of state transition if the performance improved. Performance improvement was indicated by an increase in the number of bouts and licking rate, and by a decrease in the average time of different events within a bout. Mice were then trained for at least five consecutive days on the final task (90% reward delivery, 30% chance of state transition) before the recording sessions. The reason for choosing these statistics is that they correspond to a level of environmental uncertainty that is relatively low. This allows the mice to learn the task faster than at a high level of uncertainty and to remain highly motivated during the recording sessions, thus yielding a large number of behavioral bouts.

### Electrophysiology

Recordings were made using electrode arrays with 374 recording sites (Neuropixels “Phase3A”). The Neuropixels probes were mounted on a custom 3D-printed piece attached to a stereotaxic apparatus (Kopf Instruments). Before each recording session, the shank of the probe was stained with red-fluorescent dye (DiI, ThermoFisher Vybrant V22885) to allow later track localization. Mice were habituated to the recording setup for a few days prior to the first recording session. Prior to the first recording session, mice were briefly anesthetized with isoflurane and administered a non-steroidal analgesic (Carprofen) before drilling one small craniotomy (1 mm diameter) over the secondary motor cortex. The craniotomy was cleaned with a sterile solution and covered with silicone sealant (Kwik-Sil, World Precision Instruments). Mice were allowed to recover in their home cages for several hours before the recording. After head-fixation, the silicone sealant was removed and the shank of the probe was advanced through the dura and slowly lowered to its final position. The craniotomies and the ground-pin were covered with a sterile cortex buffer. The probe was allowed to settle for 10 min to 20 min before starting recording. Recordings were acquired with SpikeGLX Neural recording system (https://billkarsh.github.io/SpikeGLX/) using the external reference setting and a gain of 500 for the AP band (300 Hz high-pass filter). Recordings were made from either hemisphere. The target location of the probe corresponded to the coordinates of the anterior lateral motor cortex, a region of the secondary motor cortex important for motor planning of licking behavior^11^. The probe simultaneously traversed the orbitofrontal cortex, directly ventral to the secondary motor cortex. In a subset of recording sessions (9 out of 11), a large portion of the probe tip ended in the olfactory cortex, ventral to the orbitofrontal cortex.

### Histology and probe localization

After the recording session, mice were deeply anesthetized with Ketamine/Xylazine and perfused with 4% paraformaldehyde. The brain was extracted and fixed for 24 hours in paraformaldehyde at 4 C, and then washed with 1% phosphate-buffered saline. The brain was sectioned at 50 μm, mounted on glass slides, and stained with 4’,6-diamidino-2-phenylindole (DAPI). Images were taken at 5x magnifications for each section using a Zeiss AxioImager at two different wavelengths (one for DAPI and one for DiI). To determine the trajectory of the probe and approximate the location of the recording sites, we used SHARP-Track ^51^, an open-source tool for analyzing electrode tracks from slice histology. First, an initial visual guess was made to find the coordinates from the Allen Mouse Brain Atlas (3D Allen CCF, http://download.alleninstitute.org/informatics-archive/current-release/mouse_ccf/annotation/) for each DiI mark along the track by comparing structural aspects of the histological slice with features in the atlas. Once the coordinates were identified, slice images were registered to the atlas using manual input and a line was fitted to the DiI track 3D coordinates. As a result, the atlas labels along the probe track were extracted and aligned to the recording sites based on their location on the shank. Finally, we also used characteristic physiological features to refine the alignment procedure (i.e, clusters of similar spike amplitude across cortical layers, low spike rate between frontal and olfactory cortical boundaries, or LFP signatures in deep olfactory areas).

### Optogenetic stimulation

To optically stimulate ChR2 expressing VGAT-expressing GABAergic interneurons we used blue light from a 473 nm laser (LRS-0473-PFF-00800-03, Laserglow Technologies, Toronto, Canada, or DHOM-M-473-200, UltraLasers, Inc., Newmarket, Canada). Light was emitted from the laser through an optical fiber patch-cord (200 μm, 0.22 NA, Doric lenses), connected to a second fiber patch-cord with a rotatory joint (FRJ 1×1, Doric lenses), which in turn was connected to the chronically implanted optic fiber cannulas (M3 connector, Doric lenses). The cannulas were inserted bilaterally inside small craniotomies performed on top of M2 (+2.5 mm anterior and ±1.5mm lateral of bregma) and barely touched the dura (as to avoid damaging superficial cortical layers). Structural glue (Super-bond C&B kit) was used to fix the fiber to the skull. The power of the laser was calibrated before every session using an optical power meter kit (Digital Console with Slim Photodiode Sensor, PM100D, Thorlabs). During the foraging task, the optical stimulation (10 ms pulses, 75 s^-1^, 5 mW) was turned on during 30% of randomly interleaved bouts. Light delivery started when the first lick was detected and was interrupted if the animal did not lick for 500 ms (which was in 98% of bouts after the last lick of the bouts).

### Statistical analysis of optogenetic manipulations

The statistical analysis of optogenetics was performed using generalized linear mixed effect models, allowing us to pool different sessions of different mice in the same model. Our N is thus the number of mice multiplied by the number of sessions and conditions (Laser OFF/ON). The different groups (control vs. inactivated) had different numbers of mice and sessions, which are reported in the results section. For each group, we fitted models with fixed effects of stimulation and random intercepts and effects of stimulation depending on mouse identity and session. For each mixed model, we report the coefficient of the fixed-effect of the stimulation predictor (Laser) ± the standard error of the estimate. We also report the p-value that corresponds to the *t*-statistic for a hypothesis test that the coefficient of the ‘Laser’ predictor is equal to 0.

To describe mixed models, we use the Wilkinson notation, with | denoting random effects. For example the formula: TimeLicking ~ 1 + Laser + (1+ Laser|Mouse) + (1 + Laser|Session), uses as predictors for the time spent licking at a foraging site a constant intercept, a coefficient for ‘laser ON’ condition that is different from ‘laser OFF’ condition, which is considered as baseline, a random intercept across mice, and a random intercept across sessions. To test the strength of the effect of stimulation on the DVs in each single session, we generated 1000 resamples of behavioral bouts in each ‘Laser OFF’ vs. ‘Laser ON’ condition, and used independent GLMs to predict the switching decision from the DVs for each resample. We compared the deviance explained of the models and the explained variance by each DV in ‘Laser OFF’ vs. ‘Laser ON’ condition, and estimated the significance of the differences. We indicated by filled dots on the plot in Fig. 3 the sessions where p-value < 0.05.

### Pre-processing neural data

Neural data were pre-processed as described previously^52^. Briefly, the neural data were first automatically spike-sorted with Kilosort2 (https://github.com/MouseLand/Kilosort) using MATLAB (MathWork, Natick, MA, USA). To remove the baseline offset of the extracellular voltage traces, the median activity of each channel was subtracted. Then, to remove artifacts, traces were “common-average referenced” by subtracting the median activity across all channels at each time point. Second, the data was manually curated using an open source neurophysiological data analysis package (Phy: https://github.com/kwikteam/phy). This step consisted in categorizing each cluster of events detected by a particular Kilosort template into a good unit or an artifact. There were several criteria to judge a cluster as noise (non-physiological waveform shape or pattern of activity across channels, spikes with inconsistent waveform shapes within the same cluster, very low-amplitude spikes, and high contamination of the refractory period). Units labeled as artifacts were discarded in further analyses. Additionally, each unit was compared to spatially neighboring units with similar waveforms to determine whether they should be merged, based on cross-correlogram features and/or drift patterns. Units passing all these criteria were labeled as good and considered to reflect the spiking activity of a single neuron. For all analyses, otherwise noted, we averaged for each neuron the number of spikes into bins by considering a 200 ms window centered around each lick. The bin-vectors were then z-scored. Because the interval between each lick was on average around 150 ms, there was little overlap between two consecutive bins and each bin typically contained the number of spikes associated with only one lick.

### Predicting choice from DVs

All data analyses were performed with custom-written software using MATLAB. We used logistic regression^53^ to estimate how DVs predicted the choice of the animal (i.e., the probability that the current lick is the last in the bout). Using Glmnet for Matlab (Qian, J., Hastie, T., Friedman, J., Tibshirani, R. and Simon, N., 2013; http://www.stanford.edu/~hastie/glmnet_matlab/) with binomial distribution, model fits were performed with DVs as predictors. We used 5-fold nested cross-validation and elastic net regularization (*α* = 0.5). To assess a metric of model fit, we calculated the deviance explained (as implemented by the devianceTest function in Matlab). The deviance explained is a global measure of fit that is a generalization of the determination coefficient (r-squared) for generalized linear models. It is calculated as:

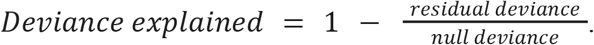

The residual deviance is defined as twice the difference between the log-likelihoods of the perfect fit (i.e., the saturated model) and the fitted model. The null deviance is the residual deviance of the worst fit (i.e the model that only contains an intercept). The log-likelihood of the fitted model is always smaller than the log-likelihood of the saturated model, and always larger than the log-likelihood of the null model. As a consequence, if the fitted model does better than the null model at predicting choice, the resulting deviance explained should be between 0 and 1. When the fitted model does not predict much better than the null model, the deviance explained is close to zero.

### Simulated behavior sessions

To test the logistic regression model, we simulated behavioral sessions of an agent making decisions using a logistic function and the DV of the inference strategy (consecutive failures). For each simulated session, the slope and the intercept of the logistic regression in the ground truth model were chosen to fit the distribution of the total number of licks in each bout from the real data. To estimate the parameters of the ground truth model (slope and intercept), we then fit a logistic regression model to predict the leaving decisions of this simulated agent using the consecutive failures DVs.

### Predicting DVs from neural population

We used a generalized linear regression model for Poisson response^54^ to predict each DV given the activity of the neural population (or facial motion, or both). Specifically, we predicted the DV *A* given the neural activity *x*, by learning a model with parameters, *β*, such as *A* = *exp*(*β*_0_ + *βx*). The Poisson regression with log-link is appropriate to model count data like the DVs studied here. To enforce positivity of the count responses, we shifted all the DVs to have a minimum value of one. Model fits were performed on each session separately. We employed elastic net regularization with parameter α = 0.5. Additionally, we performed a cross-validation implemented by cvglmnet using the lambda_min option to select the hyper-parameter that minimizes prediction error. To assess the predictive power of the model, we also implemented a nested cross-validation. Specifically, the model coefficients and hyperparameters were sequentially fit using a training set consisting of four-fifths of the data and the prediction was evaluated on the testing set consisting of the remaining one-fifth. The method was implemented until all the data had been used both for training and testing. The deviance explained reported as a metric of the goodness of fit was calculated from the cross-validated results. The final *β* coefficients were estimated using the full dataset.

### Comparison between brain regions

To ensure fair comparison between brain regions with different numbers of recorded neurons, we excluded regions with very low numbers of recorded neurons (i.e. less than 20 neurons, n = 2 recordings in Olf excluded) and used multiple approaches to match the data from each region. One approach was to run principal component analysis of the neural data from each region and select the principal components of neural activity that predicted up to 95% of the total variance (as reported in Fig. 2). A second approach was to select a subset of the original data to match the lowest number of neurons per region in each recording (subsampling with replacement, 100 repetitions). Both approaches yielded qualitatively similar results.

### Predicting choice from neural population

We used logistic regression^53^ to estimate how the weighted sum of neural activity (i.e., the neural projections onto the weights that best predict the various DVs) predicted the probability that the current lick is the last in the bout. The model fit each recording session separately as described above using the glmnet package in MATLAB and implementing elastic net regularization with α = 0.5 and a nested 5-fold cross validation to estimate the deviance explained.

### INTEGRATE-AND-RESET model

We developed a unified theory of integration in the setting of non-sensory decision making tasks. In a wide variety of tasks, animals need to keep track of quickly evolving external quantities. Here, we considered tasks where the feedback that the animal receives is binary (e.g. reward or failure). We considered an integrator given by x_t+1_ = g(o_t+1_ = 1) • x_t_ + c(o_t+1_ = 1), if the attempt is rewarded, and x_t+1_ = g(o_t+1_ = 0) • x_t_ + c(o_t+1_ = 0), otherwise. The parameters of the integrator g(o_t+1_ = 0) and g(o_t+1_ = 1) represent the computations and are bound between zero and one (g = 1 for an accumulation, g = 0 for a reset). The parameters c(o_t+1_ = 1), c(o_t+1_ = 0) add linearly and can be negative, positive or null.

We consider different scenarios involving a combination of computations but where the optimal solution only involves a one-dimensional integration. For instance, counting tasks can be solved by a linear integration, i.e. g(o_t+1_ = 0) = g(o_t+1_ = 1)= c(o_t+1_ = 0) = c(o_t+1_ = 1) = 1, where the integrated value increases by one after each attempt regardless of the outcome. In a two-alternative forced choice and more generally in an n-armed bandit task, each arm would have an integrator that increases with rewards i.e, g(o_t+1_ = 0) = g(o_t+1_ = 1) = 1, c(o_t+1_ = 0) =0 and c(o_t+1_ = 1) = 1, and decays with failures, i.e., g(o_t+1_ = 0) = g(o_t+1_ = 1) = 1, c(o_t+1_ =0) = −1 and c(o_t+1_ = 1) =0. Even in cognitively more complex tasks, involving inference over hidden states, such as reversal tasks or foraging under uncertainty, a single integrator is often sufficient. Specifically in the foraging task studied here, the optimal solution is to integrate failures but not rewards, i.e., g(ot+1 = 0) = c(ot+1 = 0) = 1, and g(ot+1 = 1) = c(ot+1 = 1) = 0.

More generally, the model produces sequences that ramp-up with failures (i.e., g(o_t+1_ = 0) = c(o_t+1_ = 0) =1; such as the consecutive failures), and the mirror images that ramp down (i.e., g(ot+1 = 0) = 1, c(ot+1 = 0) = −1). Similarly, the model can produce sequences that ramp-up or down with rewards (i.e., g(o_t+1_ = 1) = 1, c(o_t+1_ = 1) = ±1). The model also generates sequences that accumulate one type of event and persist at a constant level with the other type (i.e., g(o_t+1_ = x) = 1, c(ot+1 = x) = ±1, g(ot+1 = y) = 1, c(ot+1 = y) = 0), such as the cumulative reward integrator or its mirror image. Finally, many sequences generated by the model (where g(ot+1 = 0) = g(o_t+1_ = 1) =0) track the outcomes (i.e., reward vs failure).

There are 36 different values that the parameters of the model can take (g(o_t+1_ = 0) and g(o_t+1_ = 1) could take the values of 0 or 1 and c(o_t+1_ = 0) and c(o_t+1_ = 1) could take the values of −1, 0 or 1). In principle, each of these defines a different model which generates a time-series when fed with sequences of binary action outcomes. The 8 of them for which c(o_t+1_ = 0) = c(o_t+1_ = 1) = 0 are trivial (constant). Of the remaining 28, not all are linearly independent. For instance, the time series generated by the model that computes ‘count’ (g(o_t+1_ = 0) = g(o_t+1_ = 1)= c(o_t+1_ = 0) = c(ot+1 = 1) = 1) is equal to the sum of the time series generated by the model that accumulates reward and is insensitive to failures (g(o_t+1_ = 0) = g(o_t+1_ = 1) = 1; c(o_t+1_ = 0) = 0; c(o_t+1_ = 1) = 1) and the time series generated by the model that accumulates failures and is insensitive to rewards (g(o_t+1_ = 0) = g(o_t+1_ = 1)= 1; c(o_t+1_ = 0) =1; c(o_t+1_ =1) =0). Thus, the rank of the space of time series is 8 (two dimensions for the linear component (*c*) of the model for each of the four possible combinations of the *g* parameters, which specify the ‘computation’ the model is performing). Out of these 8 dimensions, 4 come from models that are less interesting. Two of these are the two ‘outcome’ time series (g(o_t+1_ = 0) = g(o_t+1_ = 1) = 0), which are ‘observable’. We also only consider one time series for each of the two models, since the value of the linear component associated with the outcome that is reset makes very little difference to the overall shape of the time series. For instance, the time series generated by the two models g(o_t+1_ = 0) = 1; g(ot+1 = 1) = 0; c(ot+1 = 0) =1; c(ot+1 = 1) = 0 and g(ot+1 = 0) = 1; g(ot+1 = 1) = 0; c(ot+1 = 0) = 1; c(o_t+1_ = 1) = 1 are linearly independent but almost identical for the type of outcome sequences of interest. The remaining 4 dimensions after these ‘trival’ models are removed are spanned by the 4 basis elements that we focus on in the main text (Fig. 8). Finally, the effective dimensionality of the space of time series also depends on the temporal statistics of the outcome sequences. For the particular outcome sequences experienced by the mice (which are a function of the reward and state-transition probabilities) the effective dimensionality was low, which motivated us to focus on particular subsets of outcome sequences in Fig. 8 where the time series generated by the 4 basis elements are clearly distinct.

### LM-HMM

To test the hypothesis that animals switch between discrete decision-making strategies within single sessions, we developed a new hidden Markov model with input-driven gaussian observations modeling a time-varying linear dependence 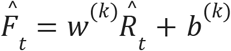 of normalized consecutive failures 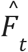 (observations) on normalized total rewards 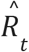 (inputs) across bouts (k) *t=1,...T*; *ϵ_t_* is i.i.d. gaussian noise with mean zero and variance σ^(k)^. For each session *m*, the normalized values 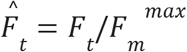 and 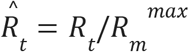 were obtained by min-maxing the raw values *F_t’_ R_t_* on their within-session max 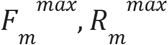. This procedure allowed us to fit a single model to all sessions where both inputs and observations were bounded between zero and one. In this LM-HMM, the slope *w*^(*k*)^, intercept *b*^(*k*)^ and noise variance *σ*^(*k*)^ depend on the hidden state *k=1*,...,*K*, each state representing a different decision-making strategy. For example, states with *w*^(*k*)^ = 0 or *w*^(*k*)^ >0 represent inference-based and stimulus-bound strategies, respectively. Large (small) values of the bias *b*^(*k*)^ represent persistent (impulsive) behavior, respectively. Other model parameters include transition probabilities between hidden states and the initial state probabilities π^(*k*)^. We fit an LM-HMM to bouts from all mice using the Expectation-Maximization (EM) algorithm to maximize the log-posterior and obtain the optimized parameters 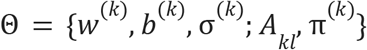. Model selection for the number of states was performed using 3-fold cross-validation by concatenating all bouts from all sessions. A model was fit to the training set, and the log-posterior of the test set was estimated (normalized by the number of bouts per test set). Because the EM may lead to local maxima of the log-posterior, for each choice of number of states, the EM algorithm was performed 5 times starting from random initial conditions. We performed model selection in two alternative ways: either by maximum likelihood estimation (MLE); or by maximum a posteriori (MAP, including gaussian prior on the weights with variance equals to 2, and Dirichlet prior on transition probabilities with alpha=2, see Ref^14^ for details on the procedure). The best number of states was chosen at the plateau of the test MLE or at the maximum of the test MAP, leading in both cases to three states. We then fit a single model to the normalized observations and inputs 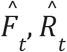, concatenating all bouts from all sessions, optimizing the model parameters Θ using MLE. Single-session values of weights and biases 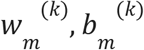 were then obtained from these normalized parameters *w*^(*k*)^, *b*^(*k*)^ as: 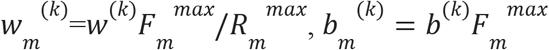.

## ACKNOWLEDGEMENTS

We thank Pietro Vertechi for insightful discussions about the project and the model and Davide Reato for support with analyses. We also thank Michael Beckert for assistance with the illustrations. This work was supported by an EMBO long-term fellowship (F.C.; ALTF 461-2016) an AXA postdoctoral fellowship (F.C.), the MEXT Grant-in-Aid for Scientific Research (19H05208, 19H05310, 19K06882, M.M.), the Takeda Science Foundation (M.M.), Fundação para a Ciência e a Tecnologia (PTDC/MED_NEU/32068/2017, M.M., Z.F.M.; and LISBOA-01-0145-FEDER-032077, A.R.), the European Research Council Advanced Grant (671251, Z.F.M.), Simons Foundation (SCGB 543011, Z.F.M.), and Champalimaud Foundation (Z.F.M., A.R.). This work was also supported by Portuguese national funds, through FCT – Fundação para a Ciência e a Tecnologia – in the context of the project UIDB/04443/2020 and by the research infrastructure CONGENTO, co-financed by Lisboa Regional Operational Programme (Lisboa2020), under the PORTUGAL 2020 Partnership Agreement, through the European Regional Development Fund (ERDF) and Fundação para a Ciência e Tecnologia (Portugal) under the projects LISBOA-01-0145-FEDER-02217 and LISBOA-01-0145-FEDER-022122.

## AUTHOR CONTRIBUTIONS

F.C. and Z.F.M. conceived the project. F.C. and M.M. designed and performed behavioral experiments. J.P.M. helped with surgery and behavioral training. F.C. designed and performed electrophysiological experiments. F.C. curated the data. F.C. and A.R. designed and performed the analyses. LM designed the LM-HMM. F.C., A.R and Z.F.M. wrote the manuscript. All authors reviewed the manuscript.

## EXTENDED DATA

**Extended Data Fig. 1:**
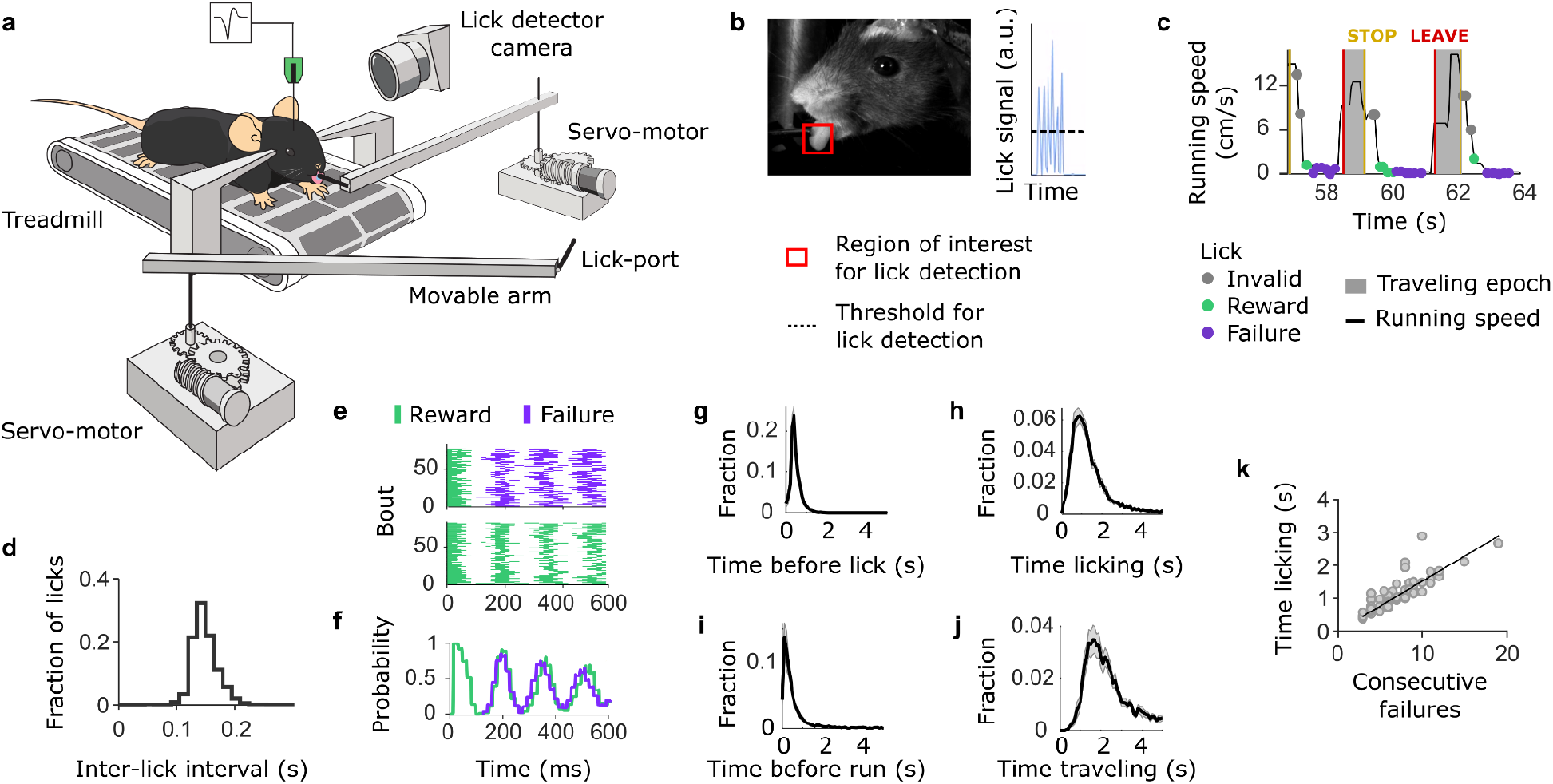
Task apparatus and behavioral properties. (a) The behavioral apparatus consists of a treadmill, coupled to two servo motors. Rotating the treadmill activates in a closed-loop fashion the movement of the arms via the motors. A mouse placed on the treadmill with its head fixed can lick at the spout from the arm in front. A camera placed on the side of the animal allows on-line video detection of the licks. (b) View from the lick detector camera. A region of interest is defined around the tongue of the animal. To detect the licks a threshold is applied to the signal within the region of interest. (c) The task consists of behavioral bouts and traveling epochs. Within a behavioral bout, the outcomes of the licks are classified into three types: reward, failure and invalid. Rewards and failures occur when the mouse slows down its running speed below an arbitrary threshold after the ‘STOP event’. The ‘STOP event’ is signaled by an auditory tone when an arm comes into place. Any lick above the running threshold is considered as invalid and always unrewarded. The traveling epoch starts after the ‘LEAVE event’, when the mouse initiates the run. (d, e, f) The licking behavior of the animals is stereotyped. (d) Histogram of the time between each lick. (e) Examples of lick raster of consecutive failures (top) and consecutive rewards (bottom). Licks are aligned at the onset of a rewarded lick and sorted based on the following events. (f) The licking frequency that corresponds to the two different examples in (e) (series of consecutive rewards in green and series of consecutive failures in purple). (g, h, i, j) Time distributions of different behavioral events (mean ± s.e.m.; n = 21 mice). The time spent licking was much greater than the time to initiate licking (between STOP event and first lick) or the time to initiate running (between the last lick and LEAVE event). Notably, engaged mice took less than half a second after the last licks to leave the site in the majority of bouts (Median time to run = 0.46 s). The running time is comparable to the licking time. (k) Monotonic relationship between the number of consecutive failures after the last reward and the time licking after the last reward (each dot represents the means across bouts for each session).

**Extended Data Fig. 2:**
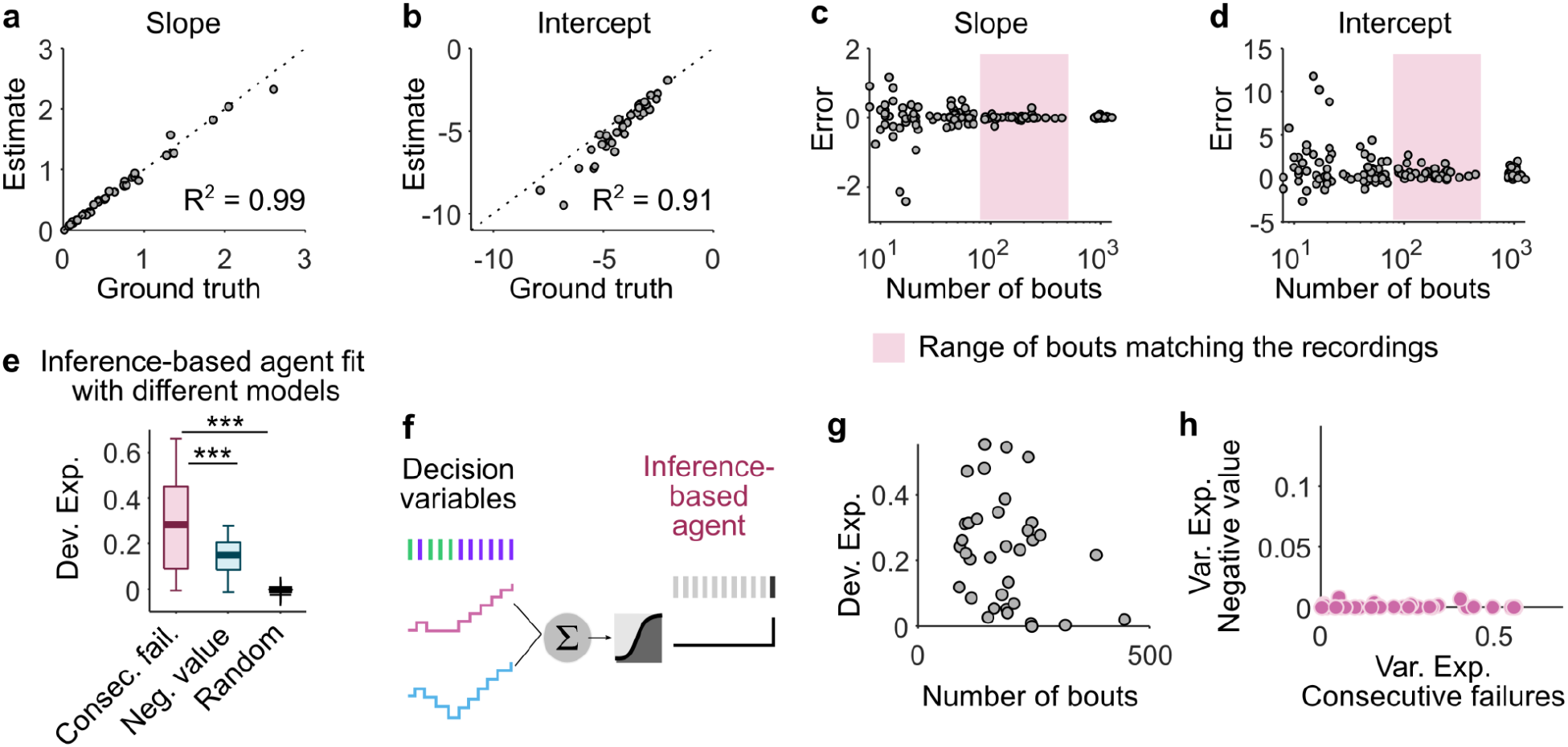
Ground truth model. (a,b) The slope (a) and intercept (b) estimates as a function of the ground truth for simulated sessions where the number of bouts matched that of real sessions. The ground truth can be recovered (R^2^ = 0.99 for the slope; R^2^ = 0.91 for the intercept) from the logistic regression. (c,d) The slope (c) and intercept (d) estimates as a function of the ground truth for simulated sessions with varying number of bouts. Overall, the ground truth can be precisely recovered for sessions with more than 100 bouts. (e) Deviance explained from a logistic regression model that fits simulated sessions of an inference-based agent using the correct model (‘Consecutive failures’), a wrong but correlated model (‘Negative value’) and a random model (where both rewards and failures are arbitrarily accumulated or reset). The deviance explained by the consecutive failures represents the upper-bound of the model performance. The deviance explained by the consecutive failures being smaller than 1 indicates that, although the ground truth can be recovered, the switching decision is not deterministic and involves some stochasticity (here the variability was matched to that of the data). However, the deviance explained by the consecutive failures is significantly greater than the deviance explained by the correlated model and the random model (Wilcoxon signed rank test, 3 stars indicate p < 10^-3^). (f) Illustration of a logistic regression model for predicting the switching decision of an inference-based simulated agent from the two different DVs (‘Consecutive failures’ and ‘Negative value’) simultaneously. (g) Deviance explained from the model in (f) as a function of the number of bouts in each session. (h) For all simulated sessions in (e), the variance explained by the ‘consecutive failures’ DV was greater than the variance explained by the ‘negative value’ DV, indicating that the model inferred the true DV.

**Extended Data Fig. 3:**
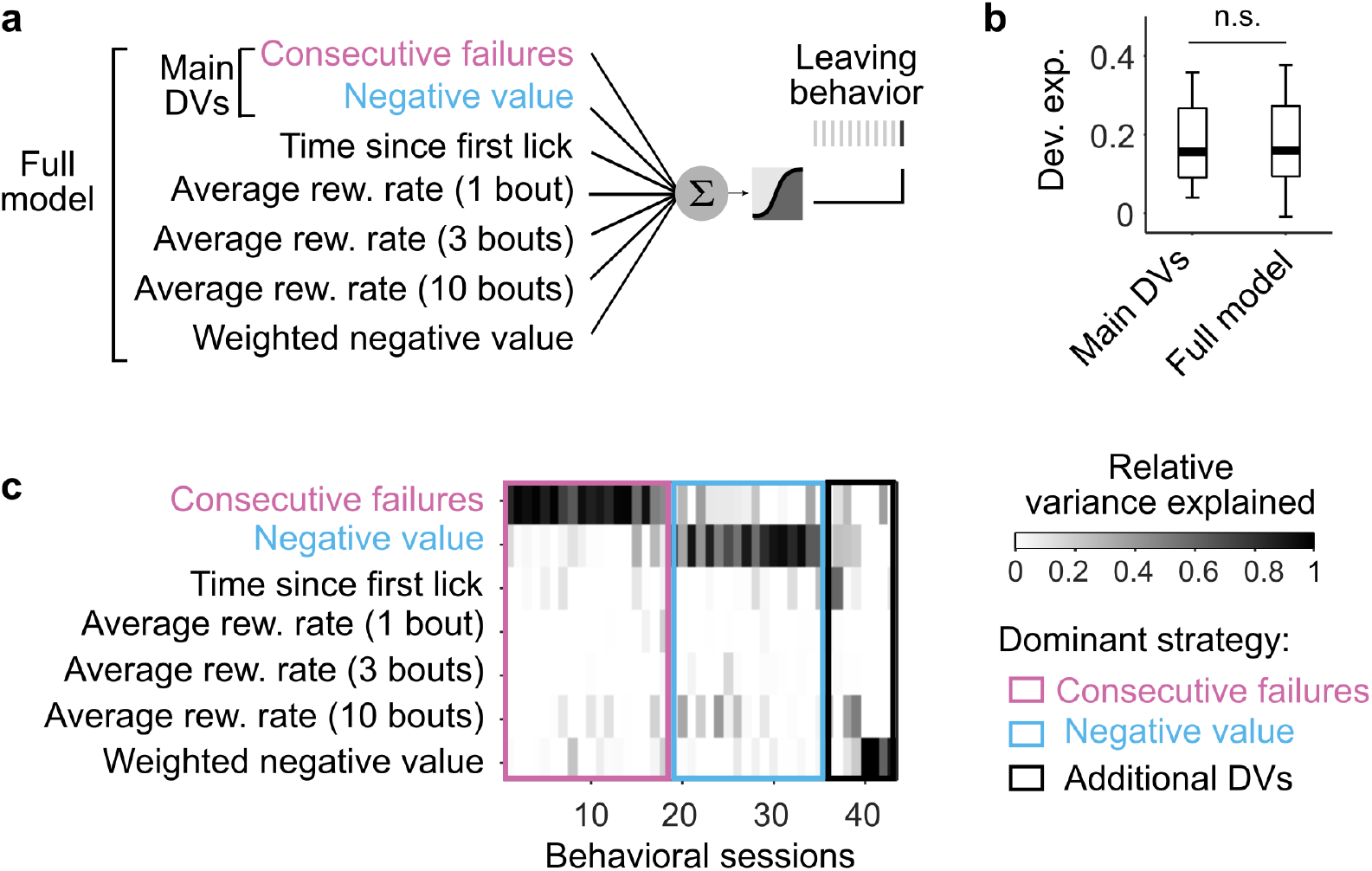
Testing alternative foraging strategies. (a) Illustration of the logistic regression model for predicting the switching decision of mice using a combination of the two main DVs, ‘Consecutive failures’ and ‘Negative value’, as well as additional DVs. Specifically, we tested 3 classes of additional DVs: 1) those relying on absolute time, 2) those relying on average reward rates, and 3) those that weigh recent evidence more strongly. The design matrix of the model thus consisted of the two main DVs, the time of each lick relative to the first lick of each bout (class 1), the average reward rate over 1, 3 and 10 previous bouts (class 2) and a version of the negative value DV that weighs recent evidence more heavily than the past ones (for class 3), such as: x_t+1_ = (1 – α)·g(o_t+1_)·x_t_ + α·c(o_t+1_), with *α* = 0.8. (b) Deviance explained from a logistic regression model that predicts choice behavior based only on the 2 main DVs (left) and from the full model that also includes the additional DVs in (a). The central mark indicates the median across behavioral sessions, and the bottom and top edges of the box indicate the 25th and 75th percentiles, respectively. The whiskers extend to the most extreme data points. There was no significant difference between the deviance explained of the two models (Wilcoxon signed rank test: p = 0.22), indicating that the additional DVs do not improve the performance of the model. (c) Relative variance explained by each predictor of the full model for each behavioral session (n = 42 sessions across 21 mice, 2 sessions per mice). The dominant DV (the one with the maximum relative variance explained) was most often the ‘Consecutive failures’ (18 sessions), followed by the ‘Negative value’ (17 sessions), and finally the additional DVs (2 session for the absolute time, 2 sessions for average reward rate, 3 sessions for the weighted negative value).

**Extended Data Fig. 4:**
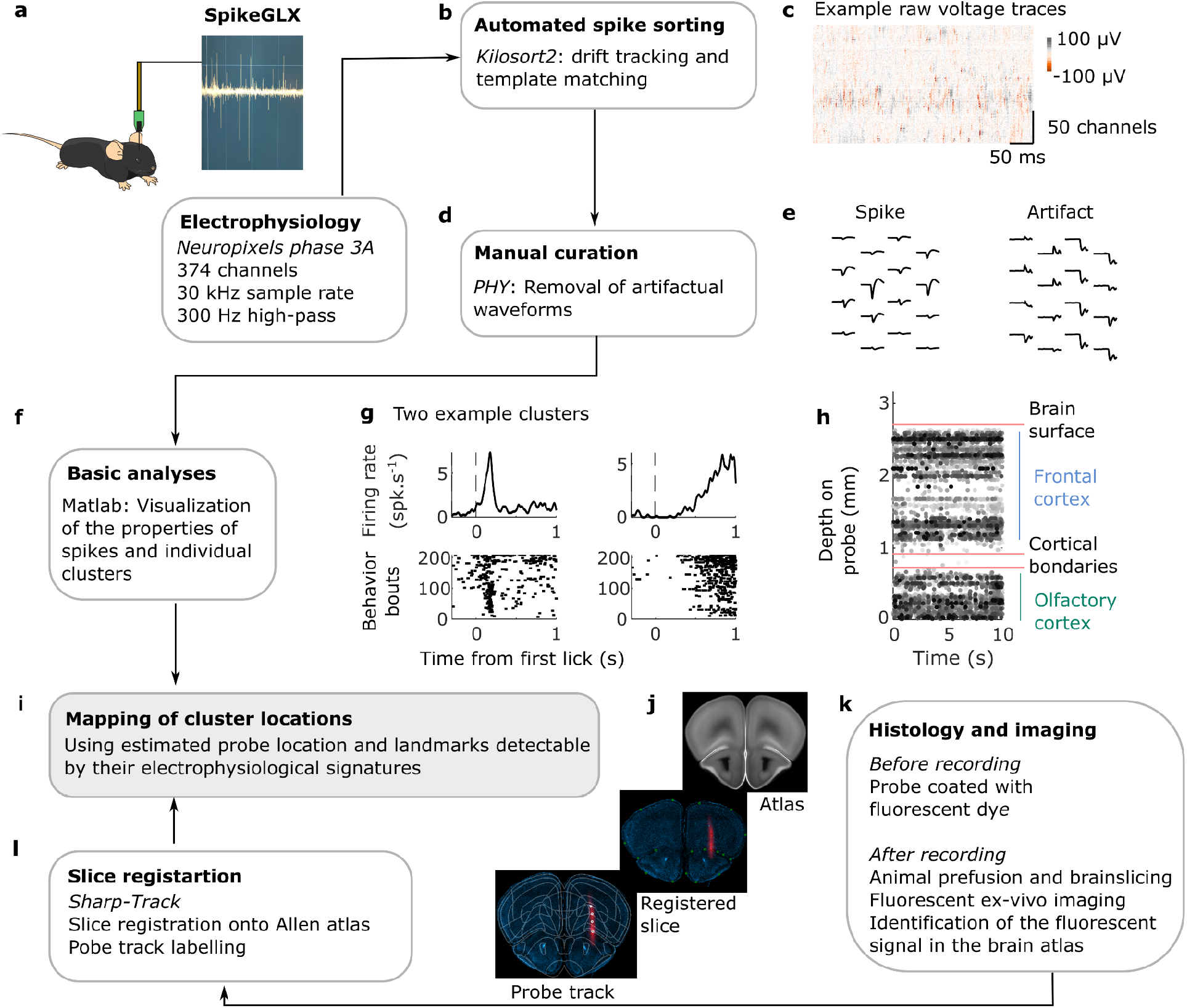
Pipeline for extracellular electrophysiology, data processing and cluster mapping. (a) Data collection from the Neuropixels probe. (b) Kilosort2 is used to automatically match spike templates to raw data. (c) Example of voltage data input to Kilosort2. Prior to the automatic sorting, the raw data is pre-processed with offset subtraction, median subtraction, and whitening steps. (d) Manual quality control is done on the outputs of Kilosort2 using PHY to remove units with non-physiological waveforms (e), contaminated refractory periods, low amplitude (less than 50 μV) or low spiking units (less than 0.5 spike·s^-1^). (f) For further quality control, visualization of peri-event spike histograms (g, top; examples histogram aligned to first lick) or scatter plots (g, bottom; example scatter plot aligned to first lick) of single neurons are made with custom-written script in MATLAB. (h, i) Example scatter plot of all neurons recorded simultaneously along the shank of the probe. This visualization helps delimitate landmarks based on electrophysiological signatures to map cluster locations. (j, k, l) Landmarks derived from electrophysiological responses are validated with estimates from histology using an open-source software (SHARP-Track).

**Extended Data Fig. 5:**
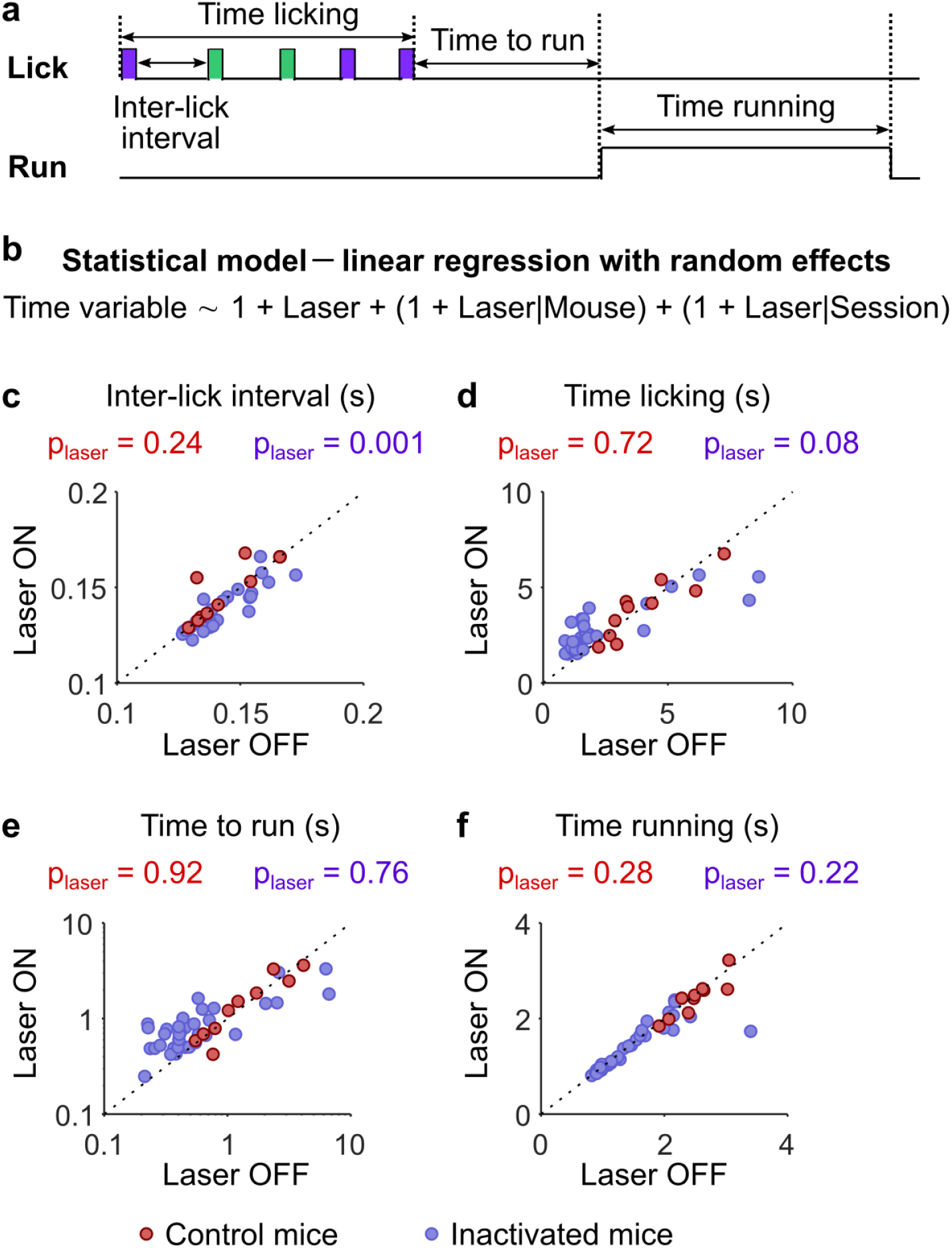
Optogenetic effect on action timing. (a) Illustration of the different action timing during a behavioral bout. (b) We used generalized linear mixed effect models to evaluate the effect of stimulation (‘Laser’ predictor) on each action timing (see Methods). The models were fit separately for inactivated and control mice (number of observations: Inactivated = 68; Control = 20). (c-f) Median timing across bouts in Laser OFF vs. Laser ON condition for each session (dots) of inactivated mice (violet) and control mice (red) mice. The p-value corresponding to the *t*-statistic for a null hypothesis test that the coefficient of the ‘Laser’ predictor is equal to 0 (p_laser_) is reported for each group of mice (color coded). (c) Fixed effect of stimulation (‘Laser’ predictor) on the inter-lick interval: Inactivated: −0.003 ± 0.0009, p = 0.001; Control: 0.005 ± 0.004, p = 0.24. (d) Fixed effect of stimulation (‘Laser’ predictor) on the time licking: Inactivated: 0.45 ± 0.26, p = 0.08; Control: −0.078 ± 0.22, p = 0.72. (e) Fixed effect of stimulation (‘Laser’ predictor) on the time to run: Inactivated: −0.075 ± 0.25, p = 0.76; Control: 0.014 ± 0.14, p = 0.92. (f) Fixed effect of stimulation (‘Laser’ predictor) on the time running: Inactivated: −0.079 ± 0.063, p = 0.22; Control: −0.061 ± 0.052, p = 0.28.

**Extended Data Fig. 6:**
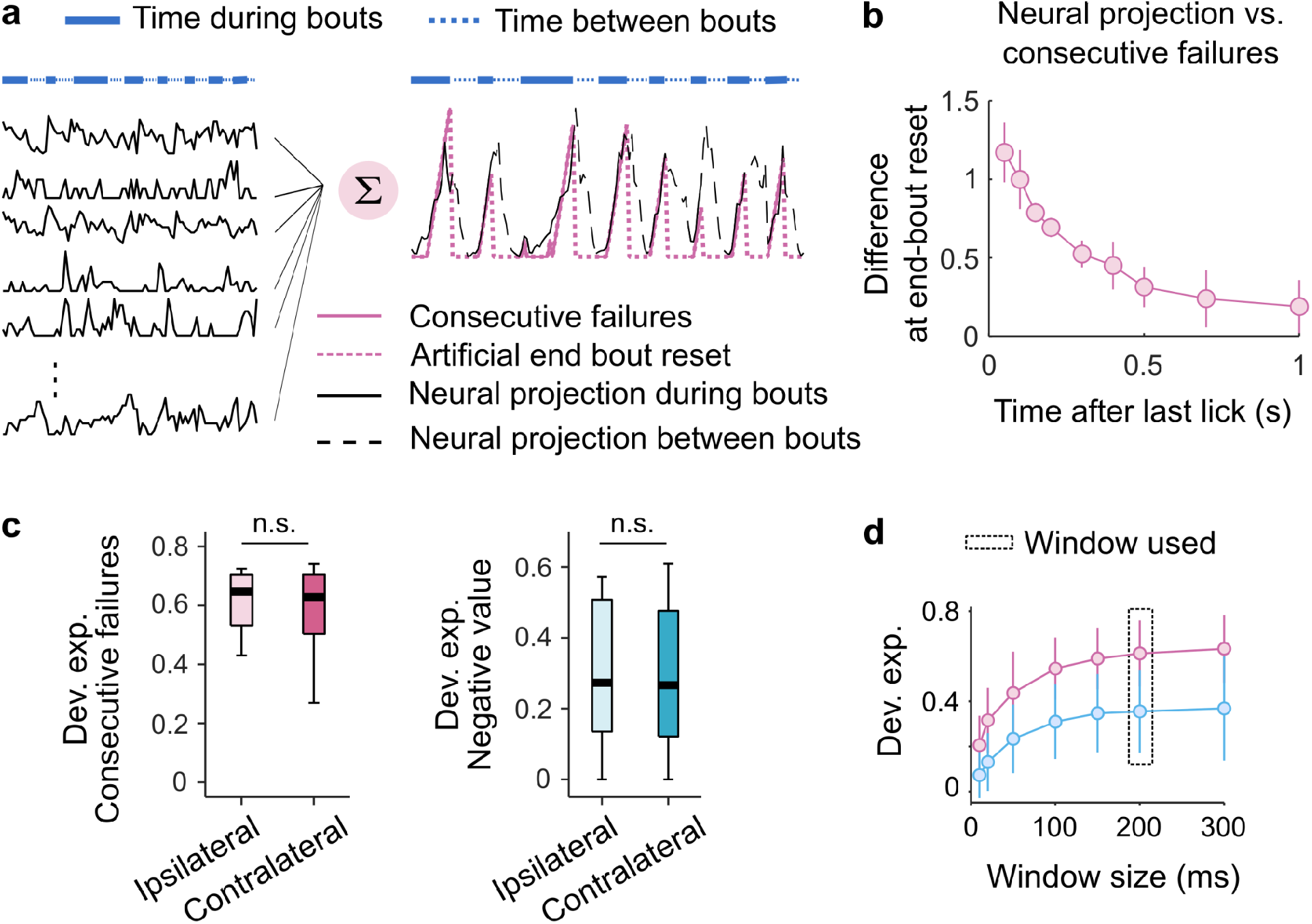
Properties of decision variables in M2. (a) Illustration of a model to estimate the time constant of the reset at the end of the bout from M2 neurons. Example consecutive failures (pink) and neural projections (black right) of the neural activity (left, example neural traces) including the activity during 2 s after the end of each bout (dashed-line). The projection of the neural activity on the decoding weights for the consecutive failure slowly ramps down until the beginning of the next bout. (b) To quantify the time constant of the reset at the end of the bout, the consecutive failures with an additional reset at the end of the bout were decoded from the neural activity. We considered the decoding projection at different times after the end of the last lick of bout ‘n’ and before the start of bout ‘n+1’ and plotted the difference between the number of the consecutive failures (dashed pink) and the neural projection (dashed black) at the end of each bout across recording sessions (median ± MAD; n = 11) as a function of the time after the last lick. The neural activity can reset at the end of the bouts with a time constant of around 200 ms. (c) Deviance explained across sessions (median ± 25th and 75th percentiles) predicted from M2 neurons for ‘Consecutive failures’ (left) and ‘Negative value’ (right) on ipsilateral vs. contralateral bouts. If the recording is performed in the right hemisphere, ipsilateral bouts are those when mice exploit the right foraging site (the right motorized arm), while contralateral bouts are those when mice exploit the left foraging site (and vice versa for recordings in the left hemisphere). We observed no significant differences in the model performance as a function of the side of the DVs (Wilcoxon signed rank test; p > 0.05). (d) This panel shows the deviance explained across sessions (median ± MAD) for DVs (Pink: ‘consecutive failures’; Blue: ‘negative value’) as a function of window sizes. In all previous analyses, the window used to count the spikes was 200 ms centered around each lick (indicated by the black rectangle), which was a good tradeoff for including a significant number of spikes while mainly considering signals related to a single lick (since the average time between each lick was around 150 ms; Fig. 4b & Extended Data Fig. 1d). Yet, a few spikes linked to the preceding or the following events could still be included in the 200 ms window, making it more difficult to evaluate the contribution of momentary evidence. Therefore, we tested whether both DVs remained decodable in M2 even when we strictly excluded all spikes from neighboring events by using smaller analysis windows. We found that the decodability of the DVs in M2 did not depend on the size of the window for widths larger than 20 ms (one-way ANOVA followed by multiple pairwise comparison tests, all p-values > 0.05 for windows size > 20 ms, both for ‘consecutive failures’ and ‘negative value’), indicating that the results are not overly sensitive to the choice of parameters.

**Extended Data Fig. 7:**
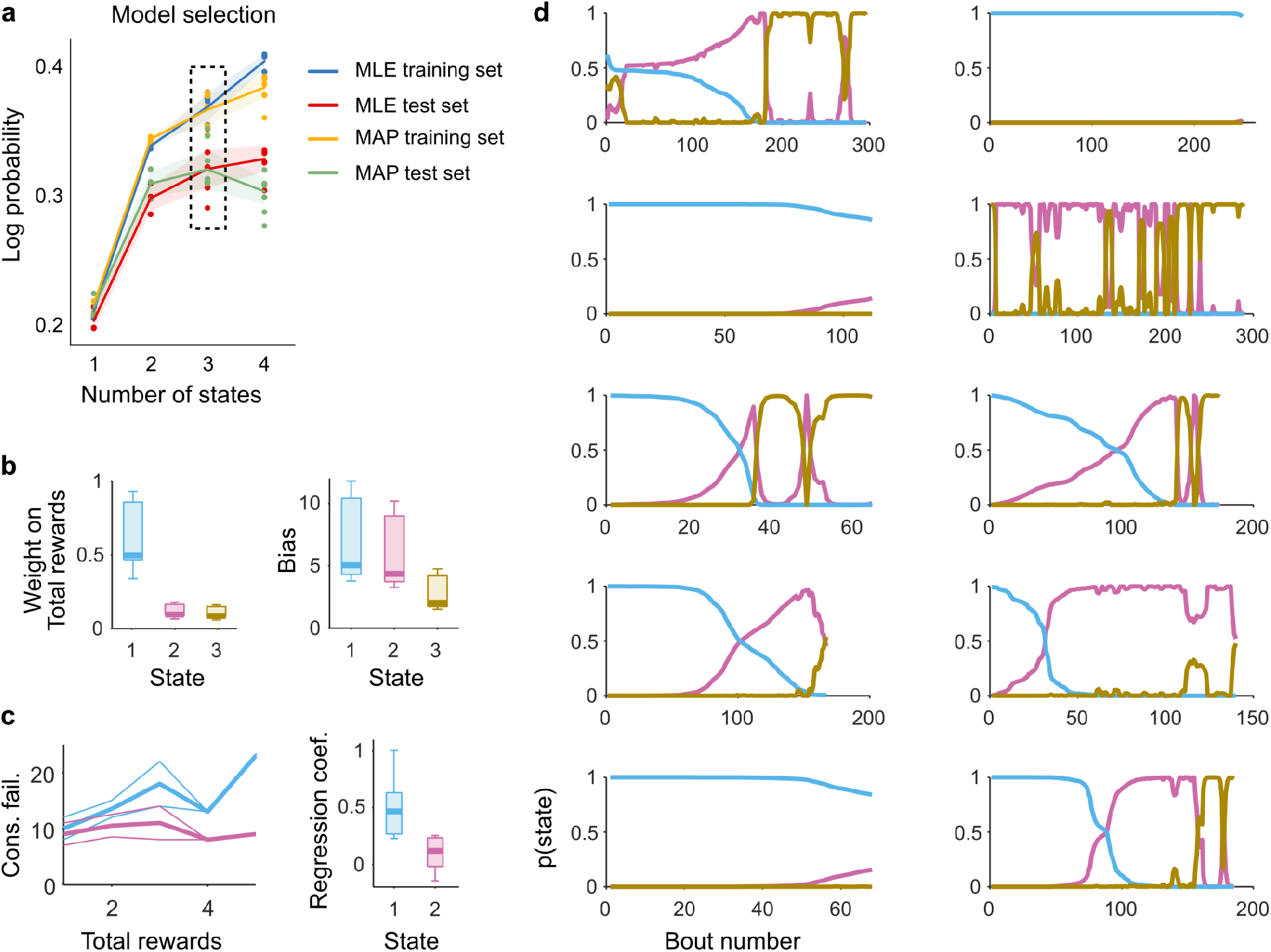
LM-HMM analysis of switch decision. (a) To determine the number of states that best capture the decision-making of mice, we fit the LM-HMM with a varying number of states and then performed model comparison using cross-validation (see Methods for details). Training and test sets log-likelihood (MLE) and maximum a posteriori (MAP, with gaussian prior on the weights and Dirichlet prior on transition probabilities) are reported in units of bits per bout. The dash-line rectangle highlights the log probability for the three-state model, which we used for all subsequent analyses. A single model was fit to all mice, where for each session the consecutive failures and prior rewards were min-maxed (i.e., divided by their max 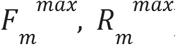), obtaining normalized weights *w*^(*k*)^ and biases *b*^(*k*)^. Single-sessions weights and biases were then obtained from these normalized parameters as 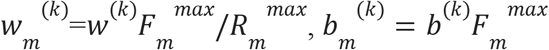. (b) Weights 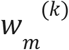 on total reward (left) and biases 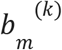 (right) across sessions *m* (median ± 25th and 75th percentiles) in the different states *k* = 1, 2, 3. (c) Consecutive failures before leaving as a function of total reward number (median ± MAD) in an example session from two different states (state 1, blue; state 2, pink). The slope coefficients of a linear regression model that predicted the number of consecutive failures before leaving as a function of the number of prior rewards in each state are shown on the right (median ± 25th and 75th percentiles across sessions). This result is consistent with the classification of stimulus-bound and inference-based strategies used in Fig. 1. (d) Posterior state probabilities for each recording session. Mice often start off the session with the stimulus-bound strategy and later switch to the inference-based strategies (in 8 out of 11 sessions).

**Extended Data Fig. 8:**
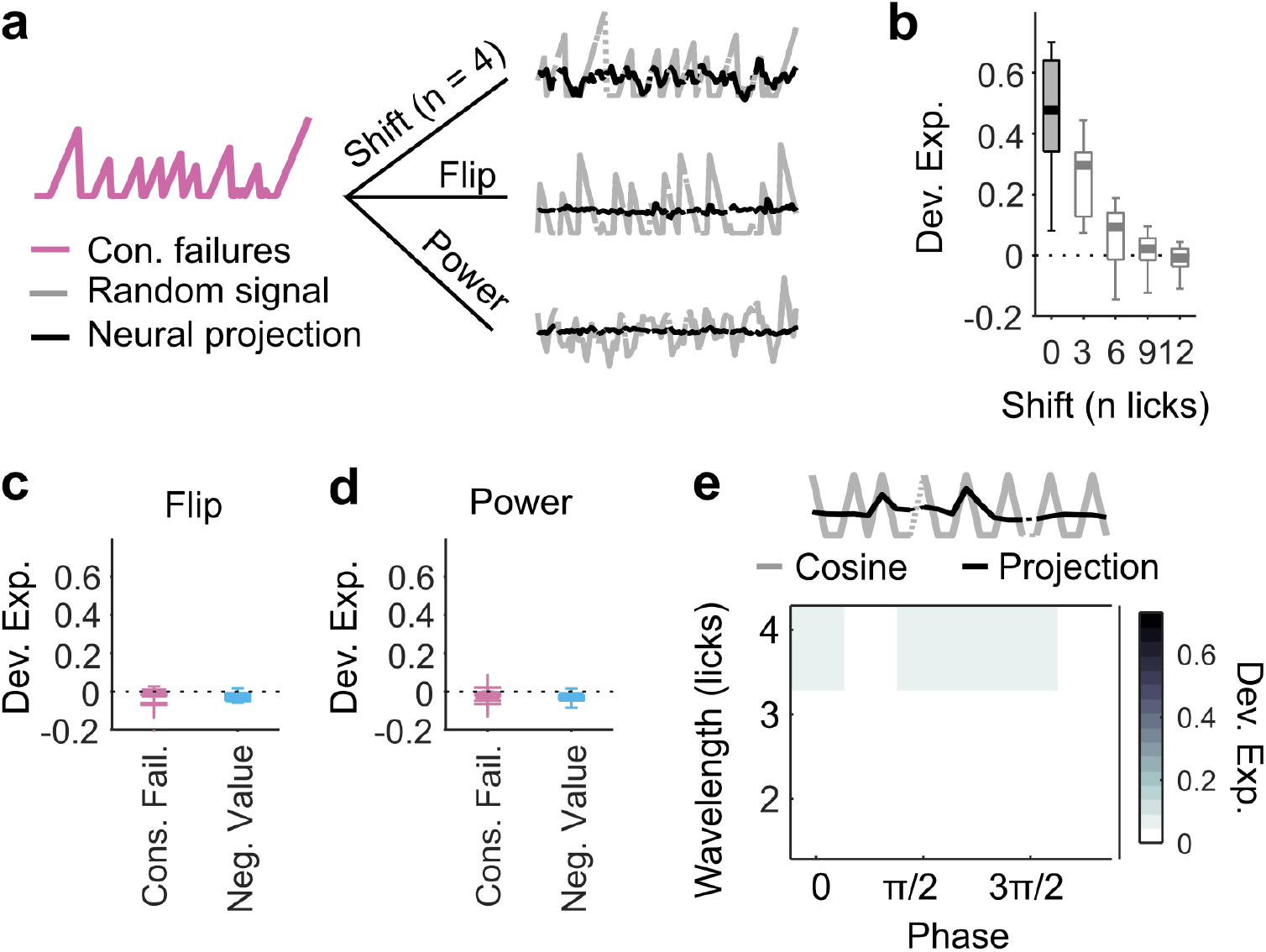
M2 does not represent arbitrary sequences. (a) A ‘near universal’ representational capacity is a feature of a computational framework known as ‘reservoir computing’ that exploits a potential functional capacity of recurrent networks to represent combinations of current inputs with previous evidence, even arbitrary ones. Thus, to test whether M2 also represented arbitrary signals, we examined whether sequences with similar temporal structure as the DVs but with no obvious relevance to the task could be decoded from M2. Here are examples of random sequences (gray) generated from one of the DVs (pink, here consecutive failures). The DV can lead to a shifted version (top right), a flipped version (middle right) or a random signal with equal power spectra. Each random signal is then decoded from M2 population activity (black traces). (b) Deviance explained (ordinate) by M2 neurons from decoding the DVs shifted by a given number of licks (abscissa). On each box, the central mark indicates the median across recording sessions, and the bottom and top edges of the box indicate the 25th and 75th percentiles, respectively. The whiskers extend to the most extreme data points. The dash black line indicates chance level (Dev. Exp. = 0). Shifting the DVs by a delay greater than their temporal autocorrelation greatly impaired their decodability (one-way ANOVA, F = 62.81, p < 10^-4^). (c) Same as in (b) but for DVs flipped across sessions. None of the flipped signals were decodable from M2 population activity. (d) Same as in (c) but for random signals with power spectra that match each DV. None of the random signals were decodable from M2 population activity. (e) Since any signal can be approximated by sums of periodic functions (Fourier analysis), we also probed the capacity of M2 to represent arbitrary temporal sequences by testing whether we could decode from M2 a basis set of cosine functions with wavelengths in the dynamic range of what we observed with integration and reset of rewards (example top gray trace; wavelength = 4 licks, phase = 0 rad). Overall, the decoding quality of the periodic function (example neural projection, top trace in black, Dev. Exp. = −0.002) was close to chance level (Dev. Exp. = 0.024 ± 0.028, median ± MAD) as seen in the matrix of deviance explained from decoding sequences with different wavelengths and phases with M2 population activity.

